# Reference genome for the highly transformable *Setaria viridis* cultivar ME034V

**DOI:** 10.1101/2020.05.02.073684

**Authors:** Peter M. Thielen, Amanda L. Pendleton, Robert A. Player, Kenneth V. Bowden, Thomas J. Lawton, Jennifer H. Wisecaver

**Author notes:** These authors contributed equally to this work (Peter M. Thielen, Amanda L. Pendleton).

## Abstract

*Setaria viridis* (green foxtail) is an important model system for improving cereal crops due to its diploid genome, ease of cultivation, and use of C_4_ photosynthesis. The *S. viridis* cultivar ME034V is exceptionally transformable, but the lack of a sequenced genome for this cultivar has limited its utility. We present a 397 Mb highly contiguous *de novo* assembly of ME034V using ultra-long nanopore sequencing technology (read N50=41kb). We estimate that this genome is largely complete based on our updated k-mer based genome size estimate of 401 Mb for *S. viridis*. Genome annotation identified 37,908 protein-coding genes and >300k repetitive elements comprising 46% of the genome. We compared the ME034V assembly with two other previously sequenced *Setaria* genomes as well as to a diversity panel of 235 *S. viridis* cultivars. We found the genome assemblies to be largely syntenic, but numerous unique polymorphic structural variants were discovered. Several ME034V deletions may be associated with recent retrotransposition of *copia* and *gypsy* LTR repeat families, as evidenced by their low genotype frequencies in the sampled population. Lastly, we performed a phylogenomic analysis to identify gene families that have expanded in *Setaria*, including those involved in specialized metabolism and plant defense response. The high continuity of the ME034V genome assembly validates the utility of ultra-long DNA sequencing to improve genetic resources for emerging model organisms. Structural variation present in *Setaria* illustrates the importance of obtaining the proper genome reference for genetic experiments. Thus, we anticipate that the ME034V genome will be of significant utility for the *Setaria* research community.

## INTRODUCTION

Grasses of the genus *Setaria* represent diverse species, with phenotypes ranging from the domesticated food crop foxtail millet, *S. italica*, to its weedy ancestral progenitor, green foxtail, *S. viridis* (Li and Brutnell 2011). Simple growth requirements, small stature, and short lifecycle make *Setaria* a tractable monocot model system for studying C_4_ photosynthesis (Brutnell *et al.* 2010; Li and Brutnell 2011; Van Eck and Swartwood 2015). Furthermore, close phylogenetic relationships with agriculturally important crops such as maize and sorghum promise to inform genetic and cell biology knowledge of other food crops of global importance. Current genome resources for *Setaria* include a reference genome for *S. italica* (Bennetzen *et al.* 2012; Zhang *et al.* 2012) based on the cultivar Yugu1, a variety of foxtail millet widely grown as a food crop in China. Additionally, a *de novo* assembly of *S. viridis* cultivar A10.1 (hereafter referred to as A10) was recently made available alongside low coverage resequencing of more than 600 *Setaria* ecotypes (Huang *et al.* 2019).

Efficient genetic modification is a primary requirement for development of any model organism. Approaches for *Setaria* protoplasting, particle bombardment, and *Agrobacterium tumefaciens* infection have been demonstrated (Brutnell *et al.* 2010; Van Eck *et al.* 2017; Mookkan 2018; Van Eck 2018). Historically, much of the *Setaria* literature referred to A10 for phenotypic evaluation, which made selecting it for whole genome sequencing a logical choice. However, many laboratories have opted to use the *S. viridis* cultivar ME034V (hereafter referred to as ME034V) due to its exceptionally high transformation efficiency, informally described at greater than 80% infected calli giving rise to at least one independent transgenic line (Acharya *et al.* 2017; Zhu *et al.* 2017). Given its high transformability and phenotypic similarity to A10, transition to ME034V as the cultivar of choice for research involving genetic modification offers significant advantages by reducing the resources required for time consuming and highly technical transformation protocols.

In this study, we report a new *de novo* genome assembly for ME034V. This genome was generated using a multi-step assembly approach, with overlap and layout performed using ultra-long Oxford Nanopore Technologies (ONT) reads (read N50 = 41 kb) and consensus polishing with Illumina sequencing. This assembly captures 397.03 Mb of total sequence, represented in just 44 contigs (contig N50 = 19.5 Mb). Additionally, we provide an updated average *S. viridis* genome size estimate of 401 Mb based on k-mer representation of multiple *S. viridis* accessions. If accurate, this estimate suggests that our new assembly, as well as previous genome assemblies, capture approximately 99% of the total genetic content of green foxtail. Included with this genome release is the *de novo* annotation of 37,908 genes, of which greater than 96% were assigned to orthologous gene families with other grasses and 60% were assigned a functional annotation. The ME034V genome assembly is highly syntenic with the two other *Setaria* genomes that are publicly available (Zhang *et al.* 2012; Huang *et al.* 2019). Our long-read sequencing provides increased resolution in regions of high repeat content, allowing for the discovery of novel insertions and other structural variants. We extended this analysis to short read alignments from over 200 additional *Setaria* cultivars to identify a dataset of 421 polymorphic structural variants, many of which contained transposable elements (TEs). Our analysis indicates that particular repeat families (*e.g. copia, gyspy*, and MULE families) show recent retrotransposition potential in *Setaria*. Taken together, the results of our study highlight genome variation between closely related cultivars of the same species and will be a valuable genetic resource for the research community that takes advantage of the uniquely high transformation efficiency of ME034V.

## MATERIALS AND METHODS

### Genome size estimation

Unprocessed Illumina sequencing data was acquired from the SRA for *S. italica* cultivar Zhang gu and 10 *S. viridis* cultivars (Table S1). K-mer distributions were evaluated using Jellyfish v2.3.0 (Marçais and Kingsford 2011) using a k-mer size of 21, and the output was evaluated in GenomeScope v1.0.0 (Vurture *et al.* 2017) with no max k-mer coverage cutoff. Data were evaluated without processing (raw), after read quality trimming to maintain > Q20 and minimum read lengths of 35bp with cutadapt (trimmed), and after removing any reads that align to *S. viridis* chloroplast (NC_028075.1) or barley mitochondrial (AP017301.1) sequences (trimmed and filtered).

### Plant growth

*S. viridis* ME034V-1 (ME034V) seed was provided by Dr. George Jander at the Boyce Thompson Institute, and grown at JHU/APL in growth chambers (400 PAR, 31°C/22°C day/night temperatures, and 12 hour light:dark cycles). Leaf tissue was harvested two weeks post-germination, following a 48-hour dark cycle to reduce starch content. Upon harvest, tissue was stored at -80°C until DNA extraction.

### DNA sequencing

DNA isolation was performed with frozen leaf tissue that was disrupted in a liquid nitrogen cooled mortar and pestle, and incubated in CTAB buffer with 20ug/mL proteinase K at 55°C for 30 minutes prior to purification with one round of chloroform and two subsequent rounds of phenol:chloroform:isoamyl alcohol.

ONT sequencing on the MinION platform utilized a ligation-based motor protein attachment approach (SQK-LSK108) and R9.4.1 flowcells (FLO-MIN106) using the protocol “1D gDNA long reads without BluePippin” (version GLRE_9052_v108_revD_19Dec2017). This protocol incorporates treatment of input DNA with NEBNext FFPE DNA Repair Mix (M6630) prior to end repair, A-tailing, and subsequent ligation of the Oxford Nanopore motor protein complex. Base calling was performed using Oxford Nanopore Guppy v2.3.5 (Wick *et al.* 2019). Reads with quality scores less than 7 were discarded.

In parallel, samples were sequenced on the Illumina NextSeq platform using Nextera XT library preparation reagents and v2 2×150bp paired-end sequencing reagents. Both ONT and Illumina sequencing libraries were generated from the same source material, and reads are available for download at the NCBI Sequence Read Archive (NCBI: PRJNA560942). Additional data for ME034V was downloaded from the NCBI SRA (NCBI: SRR1587768).

### Genome assembly

An overlapping read file from ONT data was generated using minimap2 v2.15-r911-dirty (Li 2016). These super-contiguous sequences and the original input read file were then assembled using miniasm v0.3-r179 (Li 2016). In order to find the best initial assembly for polishing and scaffolding, a range of miniasm parameter combinations were executed with each resulting assembly was evaluated for total contig count and length. Specifically, a four-feature parameter space for miniasm was explored, yielding 54 unique parameter set tests (Figure 1). 1) Minimum match lengths [-m] 25, 50, and 100 were evaluated with 100 selected as the optimal parameter. 2) Minimum overlap identifies [-i] 0.01, 0.03 and 0.05 were evaluated with 0.05 selected as the optimal parameter. 3) Minimum spans [-s] 100, 500, and 2000 were evaluated with 100 selected as the optimal parameter. 4) Maximum overhang lengths [-h] 1e4 and 1e5 were evaluated with 1e4 selected as the optimal parameter. The resulting miniasm assembly was error corrected via three rounds of polishing with ONT reads followed by 2 rounds of polishing with Illumina reads using Racon v1.4.3 (Vaser *et al.* 2017). Prior to contig polishing, the Illumina data was processed with Trimmomatic v0.35, which clipped adapters, removed bases with quality scores below Phred Q20, and removed reads less than 50 bp in length (Bolger *et al.* 2014). For SRA-acquired data, a base call quality threshold of Phred Q20 and a minimum length of 50 bp were applied using cutadapt v2.5 (Martin 2011). To annotate the four chloroplast-derived contigs, the contigs were automatically annotated using Prokka v1.14.0 (Seemann 2014), and OGDRAW v1.3.1 (Greiner *et al.* 2019) was used to generate a physical map of each annotated contig.

**Figure 1.**
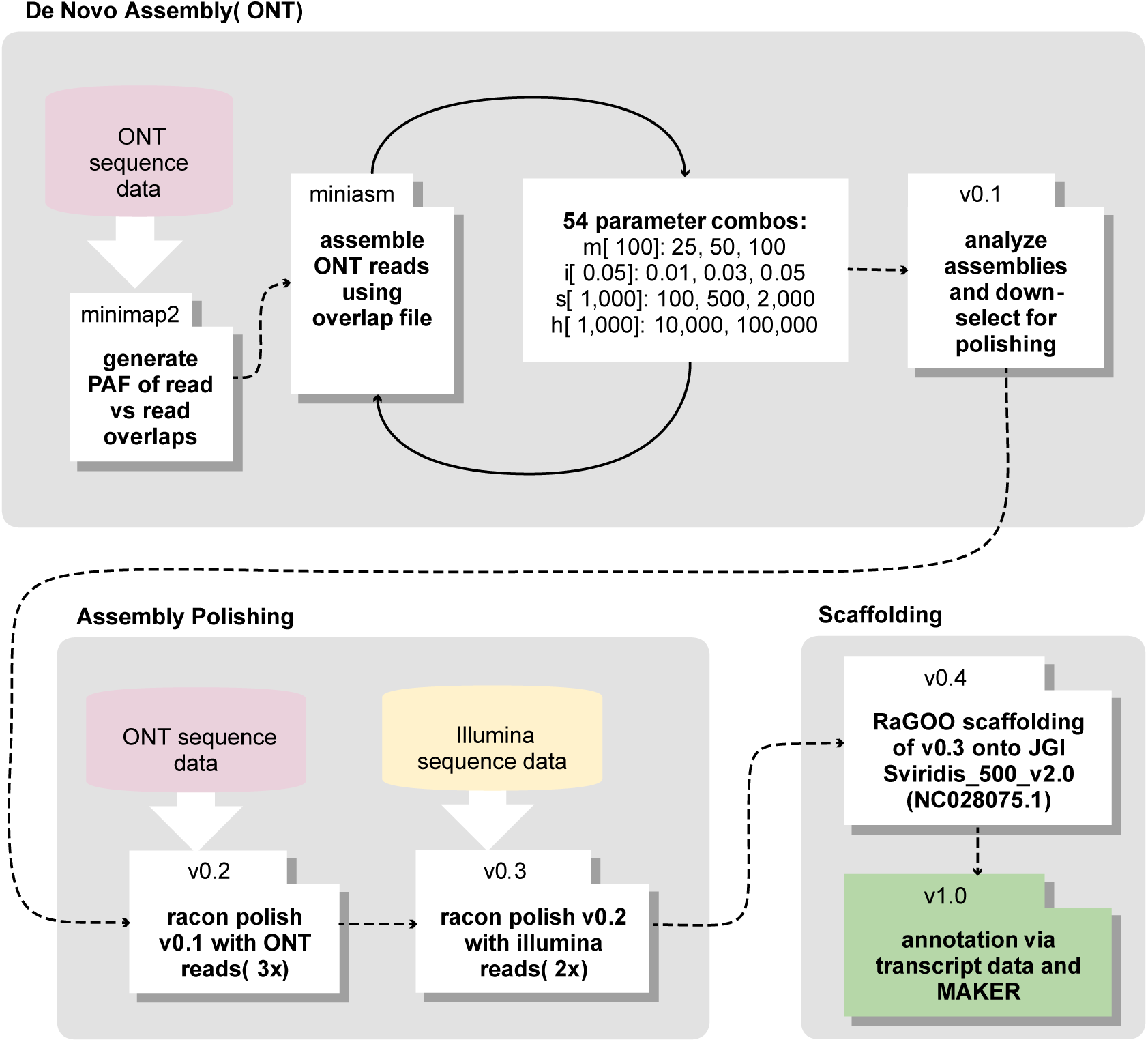
Multistage assembly pipeline. ONT long-reads were assembled *de novo* to generate assembly v0.1. The resulting assembly was first polished with ONT reads and then with Illumina short-reads to create assemblies v0.2 and v0.3, respectively. The assembly contains were scaffolded to chromosome-level pseudo chromosomes to generate the final assembly v1.0

To orient the contigs into chromosome-level scaffolds, the fast and accurate reference-guided scaffolding tool RaGOO v1.1 (Alonge *et al.* 2019) was used to scaffold the polished contigs on to Joint Genome Institute’s A10 genome assembly (NCBI: GCA_005286985.1; JGI Sviridis_500_v2.0) and the *S. viridis* chloroplast DNA sequence (NCBI: NC028075.1). Assembly statistics were calculated using QUAST-LG v5.0.2 (Gurevich *et al.* 2013). Lastly, assembly gaps in the ME034V genome were manually inserted as 100 Ns between contigs already anchored to the A10 assembly in a reference-guided manner.

### RNA sequencing and processing

Total RNA was extracted from leaf, root, and shoot tissue from four ME034V plants using the Promega SV total RNA isolation kit. RNA-seq libraries were constructed and sequenced at the Purdue Genomics Core Facility at Purdue University using Illumina’s ribominus TruSeq Stranded Total RNA library prep kit. The libraries were sequenced on the Illumina NovaSeq S4 platform (2×150), resulting in approximately 700 Gb of raw read sequence. Initial quality filtration of FASTQ with Trimmomatic v0.36 (Bolger *et al.* 2014) removed reads less than 100 bp in length (MINLEN:100), Illumina TruSeq adapters (ILLUMINACLIP:TruSeq_PEall.fa:2:30:10), performed sliding window quality filtrations (SLIDINGWINDOW:4:20), and required average total read quality scores to be at least 20 (AVGQUAL:20). Reads were merged across all samples and error corrected using tadpole (BBTools v37.93) (Bushnell *et al.* 2017) using a kmer size of 50. Error-corrected RNA-seq reads were normalized using bbnorm (BBTools v37.93) (Bushnell *et al.* 2017) to a read-depth of 25 and a minimum k-mer depth of 10 using a unique kmer size of 31. The normalized reads were aligned to the ME034V assembly with STAR v2.5.4b (Dobin *et al.* 2013) allowing alignments with an acceptable intron size range of 10-100,000 bp and a maximum multi-mapping allowance of 10. Quantification of expression was completed using Kallisto v0.45.0 (Bray *et al.* 2016). First, an index was built using the final GTF of the annotation gene set with k-mer size of 31. Raw RNA-Seq FASTQ files for twelve ME034V and eleven A10 samples (Table S2) were used as inputs for quality assessment of the gene annotation set. Following calculations of expression with Kallisto, gene expression values were calculated as the sum of transcripts per million mapped (TPM) across isoforms for each of the final primary genes (N=37,908 gene models).

### Genome annotation

*De novo* annotation of repetitive elements was conducted with RepeatModeler v1.09 (“RepeatMasker”), which also implements the *de novo* repeat finder RECON v1.08 (Bao and Eddy 2002). Nucleotide sequences of the resulting consensus repeat families were propagated using RepeatClassifier from the RepeatModeler package and used as the repeat library for RepeatMasker v4.0.7 annotation and soft masking (“RepeatMasker”). This pipeline was implemented for the ME034V assembly as well as for the *S. italica* v2 (Bennetzen *et al.* 2012), A10 v2 (Huang *et al.* 2019), *Sorghum bicolor* (Ensembl 45 release), and *Zea mays* B73 (Ensembl 45 release) genome assemblies to facilitate comparisons.

Utilizing the RNA-seq alignment file, a genome-guided *de novo* transcriptome assembly was generated by Trinity v2.5.1 (Haas *et al.* 2013) using the following parameters -- no_normalize_reads --genome_guided_max_intron 100000 --SS_lib_type RF. Gene model and protein prediction was conducted with MAKER v2.31.10 (Holt and Yandell 2011). Evidence for the first round of MAKER included the *de novo* transcript FASTA sequences from Trinity as direct EST support, the *de novo* consensus repeat calls from RepeatModeler, the FASTA sequences of the primary transcripts of A10 v2.1 (Huang et al. 2019) as alternate EST support, and alternate protein support came from over 400,000 protein sequences from closely related grass species from the Ensembl release 43 (Kersey *et al.* 2018) of *Setaria italica* (v2), *Panicum hallii* Fil (v3.1), *P. hallii* HAL (v2.1), *Oryza sativa* (IRGSP v1.0), *Zea mays* (B73 v4.0) and *Sorghum bicolor* (NCBI v3.0) and the JGI annotation of A10 v2.1 (Huang *et al.* 2019). Following the first MAKER round, Hidden Markov Model (hmm) files were generated by SNAP v2013-02-16 (Korf 2004) to inform subsequent MAKER annotations as well as training by BRAKER2 (Stanke *et al.* 2008; Hoff *et al.* 2019). The final MAKER iteration utilized SNAP, BRAKER2, all previously mentioned resources, as well as the GFF from the first MAKER round as evidence. We removed all final MAKER predicted gene models with Annotation Edit Distance (AED) scores less than 0.5. A primary gene set was determined to represent the best gene model per locus, therefore the model with the lowest AED score (best supported model per Maker) was selected. The recovery of conserved protein-coding genes was assessed using BUSCO v3 (Waterhouse *et al.* 2018) with the Eukaryota_odb9 dataset. Functional annotations were performed on the gene models using Interproscan v5.29-68.0 (Jones *et al.* 2014). Annotations of plant secondary metabolite genes was performed by hmmsearch (e-value < 1e-10; HMMER v3.1) (Eddy 2011), using hmms for 62 metabolite domains from the plantiSMASH database (“Plantismash-release”; Kautsar *et al.* 2017).

Orthologous gene families (or orthogroups) were determined using Orthofinder v2.1.2 (Emms and Kelly 2015), with sequence similar searches performed by DIAMOND (Buchfink *et al.* 2015), alignments using MAFFT v7.407 (Katoh and Standley 2013), and tree building with FastTree v2.1.7 (Price *et al.* 2010). Primary protein sets were downloaded from PLAZA monocots release 4.0 (Van Bel *et al.* 2018) for the following species: *Brachypodium distachyon, Hordeum vulgare, Triticum aestivum, Phyllostachys edulis, O. brachyantha, O. sativa ssp. Japonica, O. sativa ssp. indica, Oropetium thomaeum, Zoysia japonica ssp. nagirizaki, S. italica, S. bicolor, Z. mays, Ananas comosus, Musa acuminata, Elaeis guineensis, Phalaenopsis equestris, Spirodela polyrhiza, Zostera marina, Arabidopsis thaliana, Populus trichocarpa, Vitis vinifera, Solanum lycopersiucum, Amborella trichopoda, Selaginella moellendorffii*, and *Physcomitrella patens*. We also added the primary protein models from both A10 and ME034V to the dataset.

Orthogroups specific to ME034V were used to perform a final filtration of the gene set. We observed that many genes belonging to these orthogroups were single copy, short, and encompassed by other primary gene models. Therefore, we removed primary gene models shorter than 50 bp, as well as single-exon genes from ME034V-specific, single-copy orthogroups either without RNA-Seq expression support (TPM > 1) or mostly encompassed by another gene model (> 90% coverage). This resulted in the final gene set of 37,908 gene models.

OrthoFinder inferred duplications with branch support ≥ 0.90 were parsed from the OrthoFinder duplications.csv output. Tests for functional enrichment was performed using the PlantiSMASH hmms and the goslim_generic gene ontology (“The Gene Ontology Resource”). Hypergeometric tests were performed in python using the SciPy library *hypergeom*, and *p*-values were adjusted for multiple comparisons using the StatsModels library *multitest* with the Benjamini & Hochberg (BH) method (Benjamini and Hochberg 1995).

### Structural variation analysis

Genome-level synteny was identified using nucmer from the MUMmer package v4 (Marçais *et al.* 2018) between ME034V and A10 and *S. italica* assemblies with minimum cluster size of 65 (default, C=65) and minimum max lengths of 250bp (L=250). Dot plots were visualized using mummerplot. Smaller variations in genome alignments were identified using MUMmer’s show-diff using default parameters. Synteny plots of these small regions were generated using minimap2 v2.13 with -cx asm5 flag enabled (Li 2018) and xmatchview v1.1 (Warren 2018). Finally, read mapping support was visualized using the python-powered script package *genomeview* (“Genomeview”; Spies *et al.* 2018).

Enrichment of genome content at assembly gaps in the ME034V genome was first performed by using bedtools genomecov v2.29.1 (Quinlan 2014) to calculate the repetitive element content in non-overlapping windows across the genome. Permutation tests were completed by randomly shuffling the gap coordinates on the nuclear chromosomes 1,000 times, extracting the nucleotide sequence either 50, 100, 500, 1,000, or 1,500 base-pairs from the gap boundaries, and calculating the repeat content in these windows using bedtools. *P*-values were calculated as the proportion of permuted gaps with higher average repeat content than the observed gap content. The same process was completed using percent GC in windows surrounding known and permuted gaps.

Whole-genome sequencing data from 220 *S. viridis* and 15 *S. italica* libraries were obtained from the NCBI Sequence Read Archive (Table S3). For each library, the reads were aligned to the ME034V, A10 (v2.1), and *S. italica* (v2) genome assemblies using BWA mem v0.7.17 (Li 2013) and PCR duplicate reads were marked using Picard MarkDuplicates v2.9.0 (http://broadinstitute.github.io/picard/). The resulting BAM files were sorted using samtools v1.8 (Li and Durbin 2009) and passed to Delly v0.8.1 (Rausch *et al.* 2012) to predict inversions, tandem duplications, and deletions relative to each of the reference genomes. Delly was run with a minimum paired-end read mapping score of 20 (q=20) and a MAD insert size cutoff of 7 (s=7) for deletions. Final structural variant call sets were identified as calls with precise breakpoint support (*i.e.* split-read support), less than 5 Mb in length, passing quality scores (per Delly), mapping quality scores greater than zero, and at least five paired-reads spanning the breakpoint.

## RESULTS

### Genome size evaluation

To assess the genome size of *S. viridis*, unprocessed Illumina sequencing data were downloaded from ten *S. viridis* cultivars and one *S. italica* cultivar in the NCBI Sequence Read Archive (SRA). We performed a k-mer frequency analysis via GenomeScope using three levels of filtration: no-filtration (i.e., raw data), quality-trimmed, and quality-trimmed with removal of organellar sequences (Table S1). For ME034V, we observed a maximum haploid genome size of 432.1 Mb from raw unprocessed sequence data and a minimum haploid genome size of 391.9 Mb from quality-trimmed, organellar-filtered sequence data (Figure S1). Across all cultivars assessed, *S. viridis* cultivar ME043V produced the largest haploid genome size estimates, with a maximum of 465.6 Mb from raw unprocessed sequence data, and 421.0 Mb from quality trimmed and organellar-filtered sequence data (Table S1).

### Genome sequencing, assembly, and annotation

Sequencing data for the ME034V genome assembly consisted of 10 Gb of ONT long reads (699,624 reads with an N50 of 41 kb) and 45 Gb of 150 bp paired-end Illumina short reads. These data amount to 23-26x ONT long read coverage and 104-115x coverage with Illumina short reads assuming the maximum and minimum k-mer estimated genome size (Table S1). We then established a multistage *de novo* genome assembly workflow (Figure 1). The initial assembly was performed using minimap2 and miniasm with parameter settings optimized for long, noisy ONT reads (Li 2016). In order to find the best initial assembly for polishing and scaffolding, a range of miniasm parameter combinations were executed, and each resulting assembly was evaluated for total contig count and length (assembly v0.1; Figure 1). We error corrected this initial assembly via three rounds of polishing with ONT reads (assembly v0.2; Figure 1) followed by two rounds of polishing with Illumina reads (assembly v0.3; Figure 1). The resulting assembly v0.3 consisted of 48 contigs spanning 397.7 Mb, with an N50 of 19.5 Mb. Contigs were scaffolded into pseudochromosomes using the A10 (v2) reference genome to yield assembly ME034V v0.4 (Figure 1). Of the 48 total contigs, 44 correspond to A10 nuclear genome sequence (Table 1), with the remaining four contigs consisting of chloroplast genome sequence (Dataset S1). Using the chloroplast genomes of *Sorghum bicolor* and *Zea mays* as references, we determined that each chloroplast-derived contig contained a full-length chloroplast genome consisting of both short and large single copy loci and ribosomal DNA inverted repeats (Figure S2). The four chloroplast contigs are >99.9% similar and largely syntenic, with the exception of the short single copy locus in utg000045l that is inverted with respect to the other three (Figure S2). Multiple chloroplast sequences is suggestive of heteroplasmy, which has been documented in *Z. mays* and other plant chloroplasts (Oldenburg and Bendich 2004; Bendich 2007). Due to ambiguity as to the true chloroplast genome sequence, we excluded the four chloroplast-derived contigs from the final assembly (ME034V v1.0) as well as all downstream analyses.

**Table 1.** Nuclear genome assembly characteristics of ME034V v1.0.

|  | **Total Length**<br>**(bp)** | **Contig**<br>**Count** | **Contig N50**<br>**(bp)** | **Unspanned**<br>**Gaps<sup>1</sup>** | **% Masked<sup>2</sup>** |
| --- | --- | --- | --- | --- | --- |
| All | 397,031,521 | 44 | 19,521,898 | 35 | 46.02% |
| Chromosome 1 | 42,132,932 | 3 | 24,807,925 | 2 | 43.18% |
| Chromosome 2 | 48,726,069 | 3 | 22,686,334 | 2 | 43.03% |
| Chromosome 3 | 49,814,079 | 3 | 26,462,178 | 2 | 47.83% |
| Chromosome 4 | 39,642,072 | 2 | 21,223,292 | 1 | 53.00% |
| Chromosome 5 | 46,382,547 | 4 | 25,487,884 | 3 | 39.71% |
| Chromosome 6 | 36,113,639 | 6 | 11,701,730 | 5 | 54.03% |
| Chromosome 7 | 35,147,422 | 8 | 8,095,101 | 7 | 41.38% |
| Chromosome 8 | 42,437,421 | 13 | 7,531,332 | 12 | 56.60% |
| Chromosome 9 | 56,635,340 | 2 | 29,513,894 | 1 | 39.28% |

<sup>1</sup> Unspanned gaps are those without ONT read support following reference (A10) guided assignment of contiguous contigs
<sup>2</sup> Determined by RepeatMasker

Through a combination of RepeatModeler and RepeatMasker annotations, 46.02% of assembly bases were flagged as repetitive elements (Table 1; Table S4), an estimate that matches the overall *S. italica* repeat content (Zhang *et al.* 2012). As a proportion of bases masked, the most abundant classified group of mobile elements belong to the long terminal repeat (LTR) retrotransposons, constituting over one quarter of the genome (27.5%). Of these, 64,897 *gypsy-like* and 20,632 *copia-like* elements were predicted (Table S4). The ratio of approximately 3:1 *gypsy-like* to *copia-like* elements was also observed in the *S. italica* genome (Zhang *et al.* 2012). DNA transposons represented 30.8% of the repeat calls and 10.4% of assembly bases. The most abundant DNA transposon families included CMC-EnSpm (N=33,190), PIF-Harbinger (N=30,419), Tc Mariner Stowaway (N=17,178), and MULE-MuDR (N=11,521) (Table S4). To facilitate comparisons, we also processed other grass genomes through our annotation pipeline. The repeat content of sorghum and maize was 62.7% and 80.8%, respectively (Table S5). These results are consistent with the pattern that millets have less repetitive sequence than the nuclear genomes of other grasses (Haberer *et al.* 2005; McCormick *et al.* 2018).

Protein-coding genes were identified through a combination of *ab initio*, homology-based, and transcriptome-based prediction methods. A total of 37,908 gene models encoding 49,829 transcripts were predicted, with an average of 1.31 transcripts per gene (Table 2). The average protein-coding gene was 2,436 bp long and contained 4.06 exons. These numbers are comparable to the primary gene count of A10 (N=38,334) and *S. italica* (N=34,584) (Zhang *et al.* 2012; Huang *et al.* 2019). We observed that gene density was highest near the ends of chromosomes and generally replete in repeat-dense regions (Figure 2a). A notable exception was the gene sparse Chr08, which had the largest number of gaps (N=12), likely due to its high repeat content (56.6%; Table 1).

**Table 2.** Summary statistics of ME034V v1.0 primary gene models.

| Gene model statistics |  |
| --- | --- |
| No. protein-coding genes | 37,908 |
| No. transcripts | 49,829 |
| Mean gene length | 2,436 bp |
| Avg. no. exons per gene | 4.06 |
| Mean exon length | 389.04 bp |
| No. genes supported by RNA-Seq<sup>1</sup> (>1 TPM) | 23,724 (62.58%) |
| No. genes with functional annotation<sup>2</sup> | 25,628 (67.61%) |
| No. genes assigned to an orthogroup<sup>3</sup> | 36,521 (96.34%) |

<sup>1</sup> TPM > 1 from merged ME034V RNA-Seq data
<sup>2</sup> Assigned one or more Interpro or GO term

**Figure 2.**
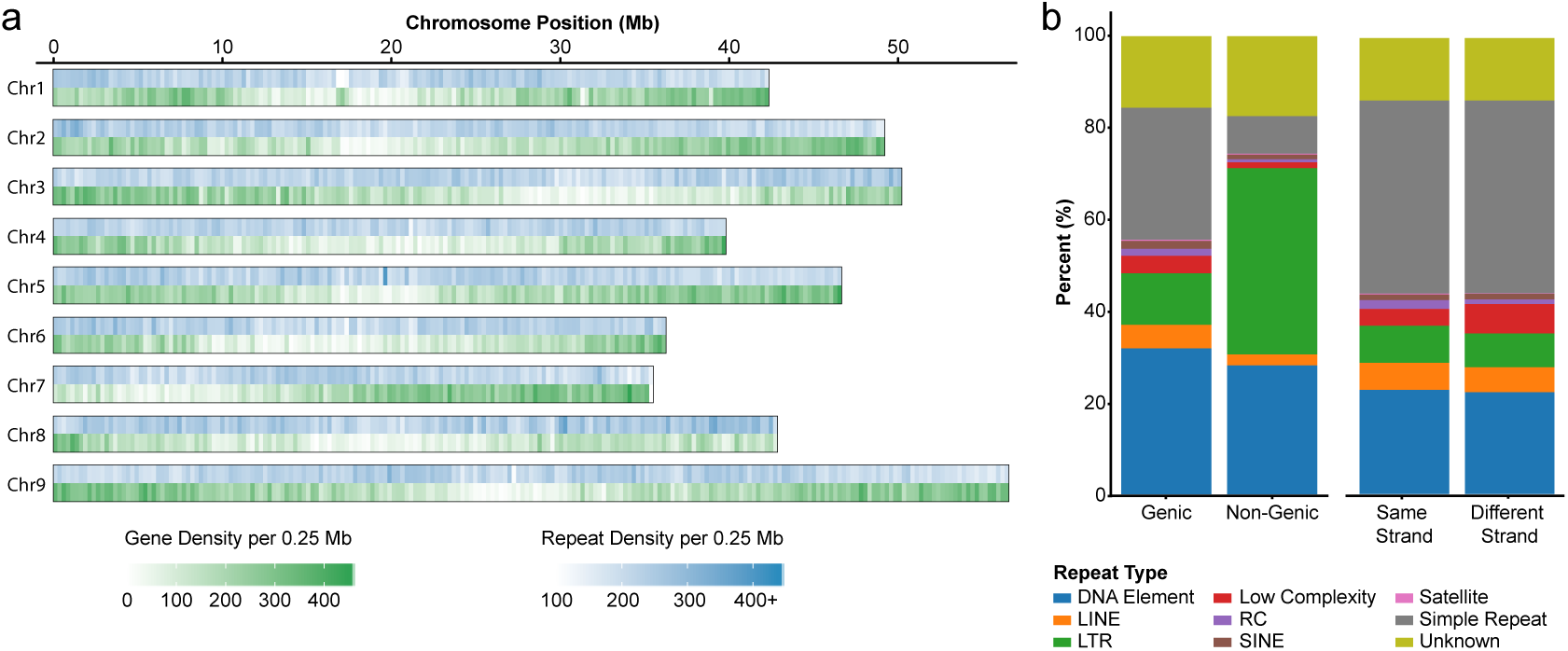
*S. viridis* ME045V genome assembly gene and repeat content. a) Gene and repeat density across the genome assembly. b) Repeat abundance by repeat type and genome location. Repeats present in genic regions are further broken up based on whether the repeat is on the same strand or different strand compared to the gene.

TEs can have profound effects on gene family evolution by altering protein-coding regions as well as gene transcriptional levels and regulation (Feschotte and Pritham 2007; Feschotte 2008). In ME034V, long interspersed nuclear elements (LINEs) make up only 1.4% of the total nuclear genome assembly; nevertheless, LINEs showed an insertion bias for genic regions, representing 5.1% of all repeats that intersect genes compared to only 2.4% of non-genic insertions (Figure 2b). In contrast, LTR elements such as *copia* and *gypsy* were nearly 3.6-fold more common in non-genic than genic space (40.7% vs 11.3%, respectively) (Figure 2b). While LINEs, LTRs, and most other complex repeat classes did not exhibit preferred insertional strand bias, rolling-circle (RC) elements (*e.g.* helitrons) were nearly twice as likely to be inserted into the same strand as a gene (Figure 2b).

Functional annotations were assigned to the majority of predicted proteins (Dataset S2). In total, 60.41% of the final 37,908 genes were assigned at least one Pfam (El-Gebali *et al.* 2019) domain, of which the majority of these domains were protein kinases PF00069 (12.06% of genes), WD40 repeats PF00400 (7.84%), and pentatricopeptide repeats PF01535 (19.34% of genes) and PF13041 (14.89%). Gene Ontology (GO) (Gene Ontology Consortium 2004) associations were also common, with 47.75% of genes assigned to at least one GO category. Additional functional annotations were assigned to 76.75%, 67.61%, 19.19%, 5.04%, 3.92% of genes using the PANTHER (Mi *et al.* 2013), InterPro (Jones *et al.* 2014), Trans Membrane (TMHMM) (Krogh *et al.* 2001), KEGG (Kanehisa *et al.* 2008), and plantiSMASH (Kautsar *et al.* 2017) databases, respectively.

### Quality assessment

To evaluate the completeness and coverage of the assembly, we aligned the Illumina gDNA and RNA reads to the ME034V genome assembly. The alignment rate of the ME034V gDNA reads was 99.14%, 98.43%, and 98.39% against the ME034V, A10, and *S. italica* genomes, respectively. We extended this assessment to samples from additional *S. viridis* and *S. italica* cultivars obtained from the SRA. Of the 235 different cultivars, 143 (60.9%) aligned best to the ME034V assembly, 36.6% aligned best to the *S. italica* assembly, and 2.56% aligned best to the A10 assembly (Table S3). Although long-read genome assemblies can exhibit collapse of repetitious or highly similar sequences (Vollger *et al.* 2019), the high alignment rate suggests minimal missing sequence and that the amount of collapsed regions in our ME034V genome assembly is not considerable.

Many of the *de novo* predicted genes in ME034V showed evidence of *in silico* expression. Over 62% of the final gene models had expression support (>1 TPM) across the merged RNA-Seq sample set from ME034V leaf, root, and shoot samples (Table 2, Table S2). Lastly, we used BUSCO (Waterhouse *et al.* 2018) to assess the completeness of the ME034V predicted proteome. Within the ME034V protein-coding gene set, 277 of 303 conserved eukaryotic genes (91.4%) were identified as complete, of those 70.6% were present in single-copy and 20.8% were duplicated.

### Comparison of assemblies

The ME034V genome assembly is largely syntenic with both A10 and *S. italica* genome assemblies, as revealed by whole genome alignments (Figure 3; Figure S3). Overall, our ME034V genome had fewer genome-specific variants when compared to A10 than to *S. italica*. However, a few large-scale variations in chromosomal structure are shared between A10 and *S. italica* that differentiate these assemblies from ME034V. These include a ∼500 kb gap on Chr01 (Figure S3a), a ∼2.5 Mb inversion on Chr02 (Figure S3b), complex rearrangements on Chr03 (Figure S3c), an inverted rearrangement on Chr05 (Figure S3e) and two inversions on Chr09 (Figure S3i).

**Figure 3.**
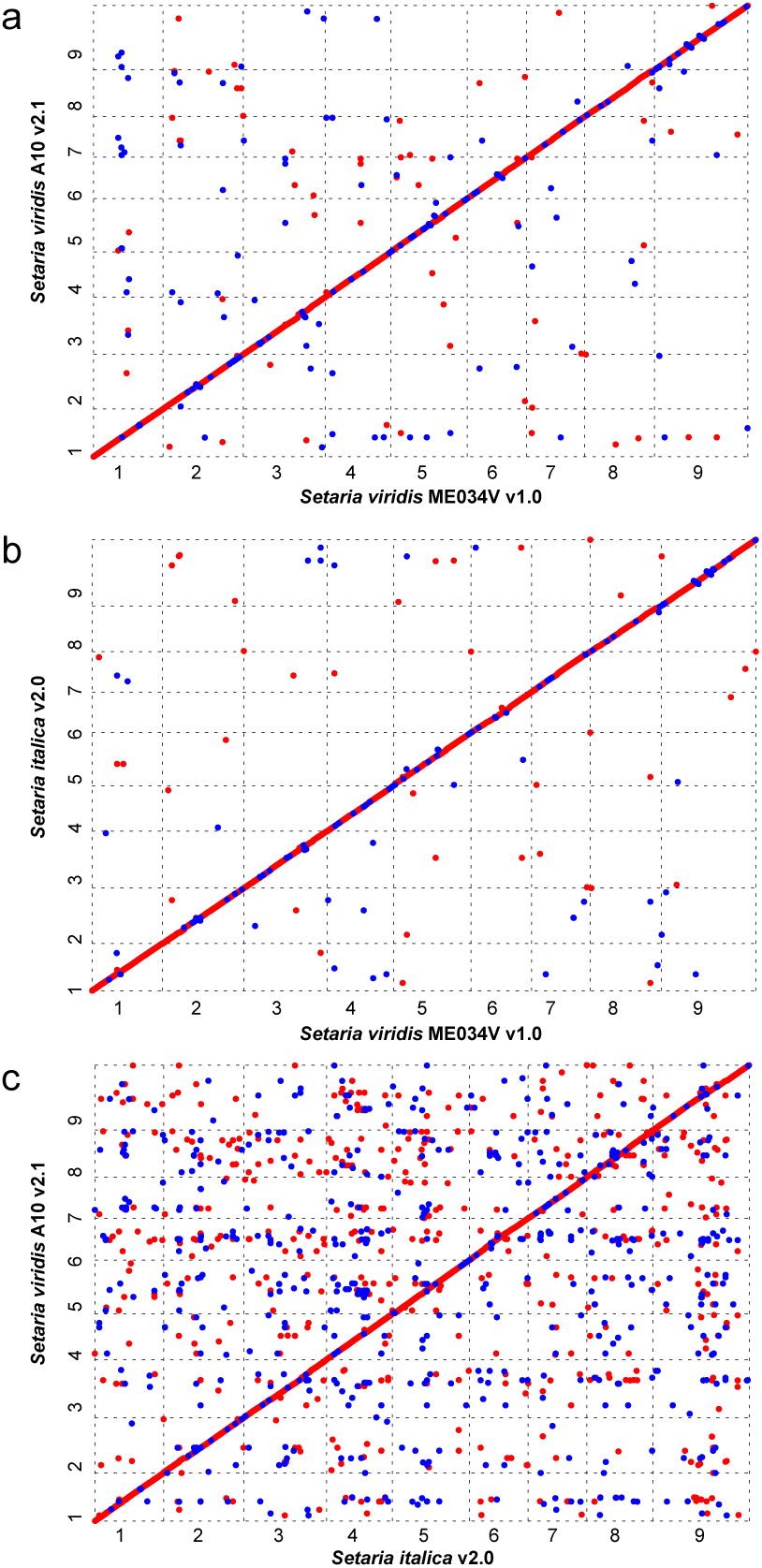
Whole genome alignments of three different *Setaria* genome assemblies. a) *S. viridis* ME034V versus *S. viridis* A10.1; b) *S. viridis* ME034V versus *S. italica;* c) *S. italica* versus *S. viridis* A10.1. Numbers along axes indicate chromosomes.

Large assembly gaps (greater than 10 bp) are far more prevalent in the *S. italica* genome (N=6,158) than both the ME034V (N=35) and the A10 (N=61) genomes. We performed a series of permutation tests to evaluate patterns of GC bias and repeat abundance in regions near gaps in our ME034V assembly. Sequences directly adjacent (50 bp flanking) to gaps in the ME034V assembly had significantly lower average GC content (*p*-value = 0.035) than randomized shuffling of sequences of similar lengths (Figure S4, Table S6). At 1,500 bp from the gap edge, the flanking sequence was significantly more likely to contain repetitive elements than expected by chance (*p*-value = 0.025). The reason significant repeat abundance was reached 1.5 kb away, rather than closer to the gap edge, is likely an artifact of annotation; if a repeat element were to extend into a gap, the truncation of the sequence could have caused repeat masking software to not call the element. Together, these data illustrate that despite the increased read-length of ONT sequences, limitations in contiguous assembly remain at genome positions of reduced nucleotide complexity and repetitive elements.

### Structural variation

Short-read alignments from over 200 *Setaria* cultivars (Table S3) against the ME034V assembly were utilized to increase resolution for identifying structural variants (SVs). This sample set is of sufficient size and genetic diversity (Huang *et al.* 2019) that even rare structural polymorphisms could be identified. Bioinformatic SV predictions are made through detection of anomalous paired read alignments. Insert sizes that are too large or too small are called as deletions or insertions, respectively, whereas read pairs with inverted orientations (*e.g.* +/+ or -/-) indicate inversions. From this analysis, we compiled a list of curated SVs less than 5 Mb in length, which contained 150 deletions, 186 duplications, and 85 inversions with read support (>5 paired-reads) at predicted variant breakpoints (Table S7). The median variant lengths for deletions was 6,630.5 bp and 2,120.0 bp for tandem duplications. The median variant length for inversions were significantly larger at 148,918 bp.

Our SV calling pipeline identified several indels containing TEs that are polymorphic within the sample population, which is suggestive of relatively recent TE activity. The most abundant *Setaria* TE class, the LTR retrotransposons *copia* and gypsy, were among these polymorphic variants. The size distribution of small deletions (<10 kb) showed several peaks that co-occurred with ME034V *copia* (peaks around 3 and 13 kb) and *gypsy* (peaks around 3 and 8kb) lengths (Figure 4). We then used synteny analyses coupled with visualization of read alignments at predicted SV breakpoints to confirm the presence of select SVs. A deletion of ME034V sequence around Chr03:33.2 Mb (DEL00053315) was predicted in five *Setaria* cultivars (Estep_ME062, Huang_TX01, Huang_MO13, Feldman MF131, Estep_ME050V; Table S7) and confirmed through breakpoint assessments (Figure S5). Synteny analyses of this region in ME034V and its homologous locus in *S. italica* (chr3:35.5 Mb) revealed that the adjacent regions are not identical in sequence but are both LTR-rich with ∼10 kb unique sequence in ME034V (Figure S6). Pairwise alignments of A10 to both ME034V and *S. italica* illustrate that the A10 assembly (Chr_03:34.8 Mb) is fully missing the locus (Figure 5a; Figure S6). Furthermore, co-linear overlaps at the apparent A10 indel is suggestive of a target site duplication (TSD) (Figure 5a), which is a signal of a retrotransposon insertion through target-primed reverse transcription (Ewing 2015). A second example includes an insertion of a *copia* element in A10 (DEL00051066) that is absent from both ME034V and *S. italica* (Figure 5b; Figure S7), which also has synteny at the putative indel breakpoint, resembling a possible TSD (Figure 5b). Read alignment patterns at the putative breakpoint indicate a homozygous insertion four (Estep_ME005, Estep_ME009, Estep_ME061V, Estep_ME059V) and heterozygous insertion for two (Feldman_MF136, Estep_ME010) *Setaria* cultivars (Figures S8; Table S7). Lastly, we identified an apparent deletion (DEL00033261) containing a *gypsy* element that is present in all three reference genomes (ME034V, A10 and *S. italica*). All three *Setaria* assemblies are syntenic at this locus (Figure 5c; Figure S9), yet three cultivars (Feldman_MF137, Feldman_MF156, and Estep_ME018) have apparent homozygous deletions based on breakpoints in their read alignments (Figure 5d and e; Figure S10).

**Figure 4.**
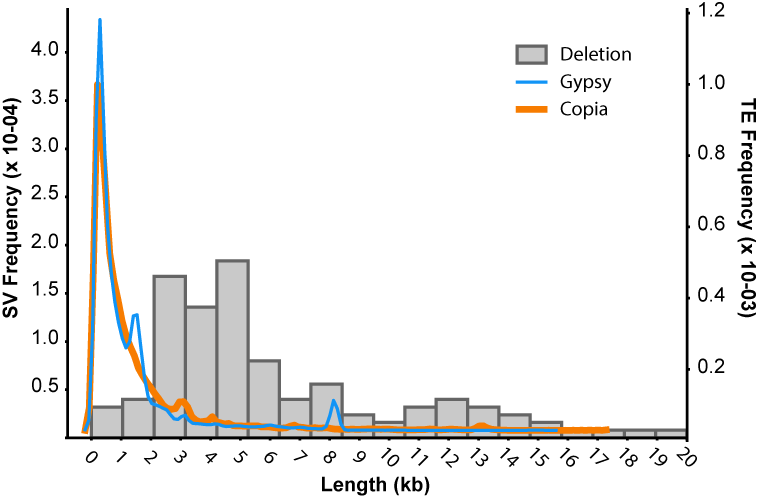
Length distribution of deletions in *ME034V* assembly compared to average size of common transposable elements. Histogram of length distributions of predicted deletions (gray bars) overlapped by density plots depicting the size distribution of annotated *copia* (orange) and *gypsy* (blue) retrotransposons in the ME034V assembly.

**Figure 5.**
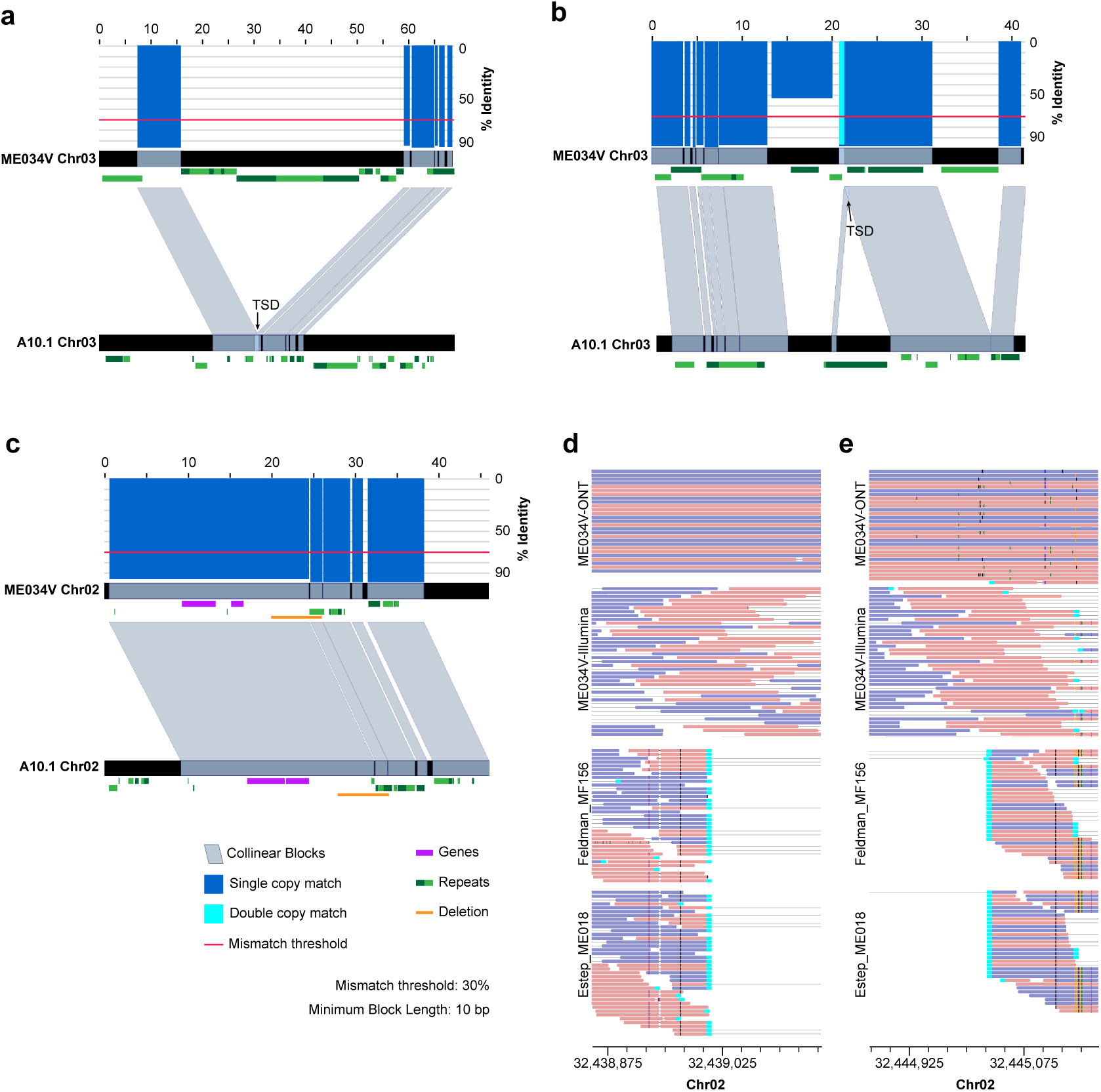
Exemplar structural variants in the ME034V genome. a) Synteny plot revealing a *copia* insertion at window Chr03:33,177,187-33,245,787 in ME034V that is missing in the homologous locus in A10.1, window Chr03:34,798,544-34,868,544 (see also Figure S5; Figure S6) b) *Copia* insertion unique to A10 (window Chr_03:28,599,689-28,645,328), and absent in ME034V (window Chr03:29,091,552-29,132,959) (see also Figure S7; Figure S8). c) Presence of a *gypsy* element shared between ME034V (window Chr02:32,419,004-32,465,022) and A10.1 (window Chr_02:31,649,962-31,696,029) is indicated by near-perfect alignment in the synteny plots (see also Figure S9; Figure S10). For all synteny plots, blue-grey bars connect the two genomes when DNA sequence with >70% identity is observed (red line indicates threshold). Blue and cyan bars above the top track indicate sequence identity along the chromosomal segment from 0-100%, with the color indicating wither single copy (blue) or double copy (cyan) matches. Purple rectangles indicate genes and green rectangles indicate LTRs (alternating hues aid in distinguishing between unique elements). The strandedness of the genes and LTRs is indicated by placing elements encoded on the forward strand higher relative to elements encoded on the reverse strand. Putative target site duplications (TSD) are indicated in collinear regions in (a) and (b). Orange rectangles in part c indicate the 1:1 homologous region absent in cultivars Estep_ME018 and Feldman_MF156; support for this deletion is visualized by split reads (cyan) at the left (d) and right (e) breakpoint of the read alignment. Read pairs are connected with gray lines, and reads on the forward and reverse strand are colored pink and purple, respectively.

### Gene family analysis

To investigate the evolution of different gene families, including those that may be unique or expanded in *Setaria* species, we performed an OrthoFinder (Emms and Kelly 2015) analysis using the protein-coding genes of ME034V and 26 other eudicot genomes, including A10 and *S. italica* (Table S8). The OrthoFinder analysis identified 28,055 unique orthogroups (predicted gene families) consisting of two or more species in the analysis (Table S9). Of the 18,973 orthogroups containing one or more ME034V sequences, 4,135 orthogroups (21.79%) were present in all species in the analysis, and 7,390 (38.95%) were found only in monocots (Figure 6a). The total number of ME034V orthogroups was comparable to the other sequenced genomes of *Setaria*, which ranged from 20,620 in A10 to 17,253 in *S. italica*. The proportion of genome-specific orthogroups was more similar between *S. italica* (6.30%) and ME034V (7.77%) than A10 (12.75%). In total, 36,521 of 37,908 ME034V proteins (96.34%) were assigned to an orthogroup containing sequence from one or more additional Poaceae (Dataset S3).

**Figure 6.**
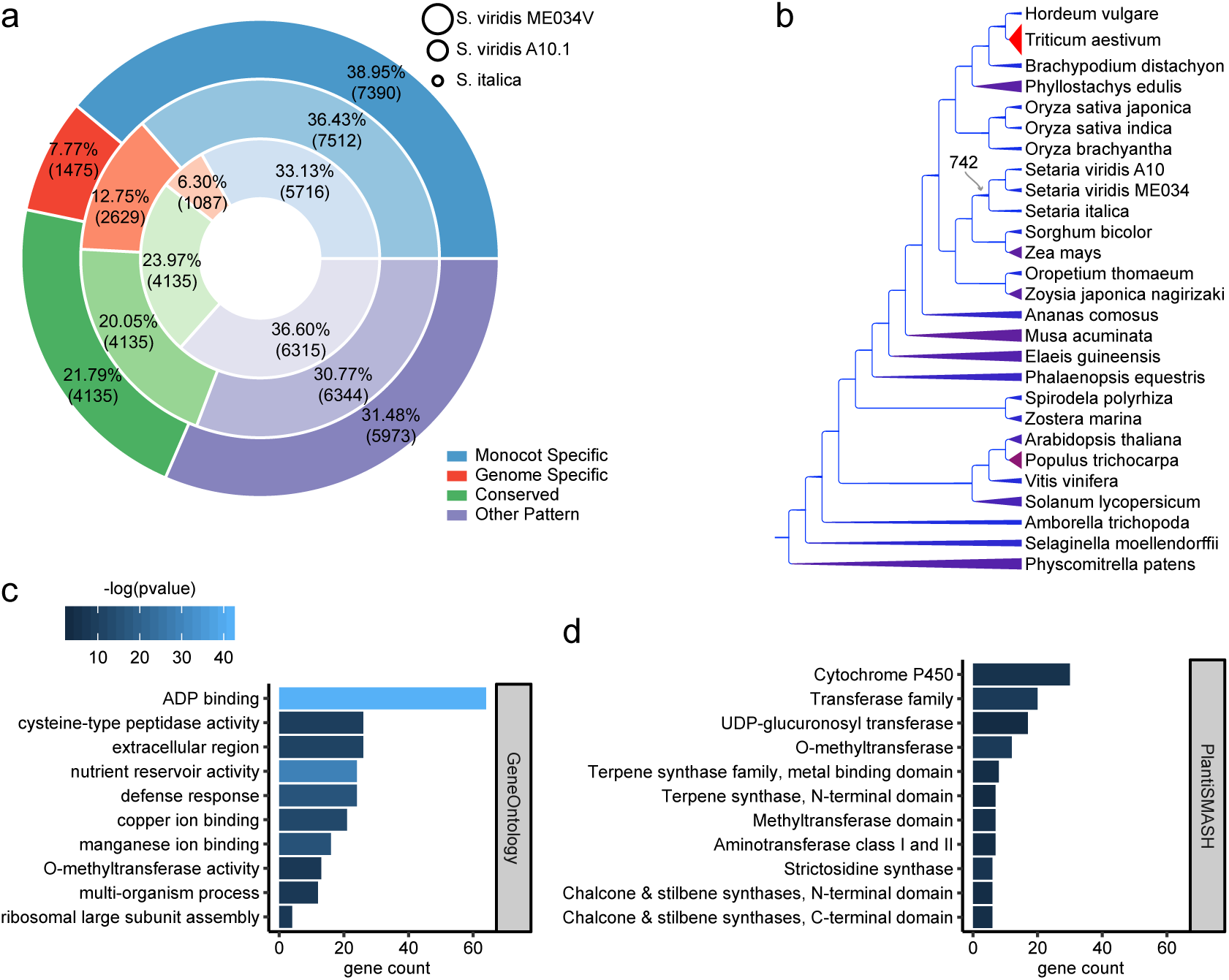
Analysis of gene families in *Setaria*. a) Comparison of orthogroups in ME034V (outer ring), A10 (middle ring), and *S. italica (inner ring).* Conserved orthrogroups (green) were present one or more times in all 27 genomes in the analysis. Monocot-specific orthogroups (blue) were present in two or more monocot genomes and absent from all others. b) The species phylogeny was taken from the PLAZA 4.0 monocots online database. Branch thicknesses and colors are scaled based on the number of predicted duplicated events to have occurred at the descendent node; thinner, blue branches indicate fewer duplications; thicker, red branches indicate more (see Table S10). The 742 duplications predicted at the *Setaria* ancestral node are indicated with the grey arrow. Tests for enrichment of functional categories was performed on this gene set: c) top ten most significantly enriched GO categories (see Table S11); d) all significantly enriched plantiSMASH specialized metabolism enzyme classes (see Table S12).

To identify orthogroups that may have expanded in the ancestor of the three *Setaria* genomes, we parsed the number of OrthoFinder predicted gene duplications at each node of the inferred species tree (Figure 6b; Table S10). The average number of orthogroups that duplicated one or more times at a given node was 1541.53 and ranged from only 2 duplications at internode 16 (the common ancestor of *Phyllostachys* and the Pooideae) to 12,238 duplications in the hexaploid *Triticum aestivum* (Figure 6b; Table S10). A total of 742 orthogroups duplicated in the last common ancestor of the three *Setaria* (Figure 6b; Table S10), and the ME034V genes that duplicated at this internode were enriched in 153 Gene Ontology (GO) categories (Benjamini-Hochberg adjusted *p-* value < 0.05; Table S11). The most significantly enriched GO categories included those for ADP and metal binding (GO:0043531; GO:0030145; GO:0005507), nutrient reservoir activity (GO:0045735), defense response (GO:0006952), extracellular region (GO:0005576), and cysteine-type peptidase activity (GO:0008234) (Figure 6c; Table S11). In addition, we checked for enrichment of enzyme families typically associated with specialized metabolic (SM) processes (Kautsar *et al.* 2017) and found that 11 SM enzyme families were enriched in the set of genes that duplicated in the ancestor of *Setaria* (Benjamini-Hochberg adjusted *p-*value < 0.05; Table S12) including those for the production of terpenes, strictosidines, chalcones, and stilbenes (Figure 6c).

## DISCUSSION

Many publications reference a genome size estimate of 490 Mb for *S. italica* and *S. viridis*, based on C-values derived from flow cytometry data (Le Thierry D’Ennequin *et al.* 1998; Bennett *et al.* 2000) and later at 485 Mb based on k-mer analysis of Illumina sequencing data (Zhang *et al.* 2012). Despite this, both *S. italica* and A10 reference assemblies are significantly smaller at 405.7 Mb and 395.7 Mb, respectively (Bennetzen *et al.* 2012; Huang *et al.* 2019). Similarly, our ME034V genome assembly totals 397 Mb.

The k-mer based genome size estimate for our assembly suggests the true *S. viridis* genome size is closer to the assembled genome sizes of ∼400 Mb. However, it is likely some repetitive regions of the ME034V genome have collapsed, as is seen in other complex genomes (Vollger *et al.* 2019). This could explain some but not all of the disagreement in genome size. Disconnect between flow cytometry and k-mer based genome size estimates has been documented by others (Pflug *et al.* 2019), and should be investigated in more detail in future analyses.

Although, the ME034V assembly is largely syntenic with the two other sequenced *Setaria* genomes from *S. italica* and A10, several SVs were identified in ME034V. Validation of many SVs by read mapping against the *Setaria* diversity panel, indicates that the SVs are unlikely to be the result of misassembly and instead represent true genome variation in the species. Identification of this genome variation, thanks in large part to the high continuity of the ME034V assembly, illustrates the utility of ultra-long DNA sequencing data to improve genetic resources for emerging model systems. Preliminary surveys of the ME034V assembly has revealed a repeat-rich landscape, with some transposable element classes displaying compelling patterns of recent mobility. Structural variant predictions in large sample sets can facilitate the identification of rare deletions or insertions. Identified from the sample set of hundreds of *Setaria* cultivars, we have presented evidence of LTR retrotransposons whose insertions are either genome-specific or completely absent in a subset of samples. Further analyses are required to validate these bioinformatic predictions in the population, assess the completeness and age of these putative TE insertions (Bennetzen *et al.* 2017), as well as evaluate their abundance in a phylogenetic context.

Lastly, our analysis of gene family evolution in *Setaria* identified hundreds (n=742) of orthogroups that likely duplicated in a common ancestor of the three genomes analyzed (ME034V, A10 and *S. italica*). These duplicated gene families appear to be enriched in processes related to specialized metabolism, nutrient acquisition, and defense response, which is consistent with previous observations that these gene families are some of the most likely to undergo frequent duplication in plants (Pichersky and Lewinsohn 2011; Chae *et al.* 2014). Future work

Altogether, our assembly of the *Setaria viridis* ME034V genome constitutes an essential resource for monocot research and further establishes *Setaria* as an ideal model plant system. Combined with the high *A. tumifericans* transformation rate of ME034V, the assembly and annotation described here will further aid in genetic manipulations, securing ME034V as the preferred *S. viridis* cultivar.

## ACKNOWLEDGMENTS

This work was supported by the DARPA Advanced Plant Technologies program to T.J.L. and J.H.W. under contract HR001118C0146. This work was supported by start-up funds from Purdue University to J.H.W., NSF Dimensions of Biodiversity Program under Grant No. DEB-1831493 to J.H.W.; this work was also supported by the USDA National Institute of Food and Agriculture Hatch Project numbers 1016057 to J.H.W..

## DATA AVAILABILITY

Raw sequencing reads used for *de novo* whole-genome assembly and the final genome have been deposited in the Sequence Read Archive database under BioProject PRJNA560942. The genome assembly has been submitted to NCBI under GenBank accession CP050795. The gene, repeat, and structural variant annotation set described in this manuscript is available for upload via a custom track hub for the University of California Santa Cruz (UCSC) Genome Browser (https://github.rcac.purdue.edu/JenniferWisecaverGroup/ME034V_Trackhub).

## SUPPLEMENTARY TABLE LEGENDS

**TABLE S1**

Estimated haploid genome length for *S. viridis* and *S. italica.*

**TABLE S2**

Sequencing library alignment rates.

**TABLE S3**

List of whole genome sequencing libraries downloaded from the NCBI SRA.

**TABLE S4**

Counts of repetitive element classifications for the nine nuclear chromosomes of ME034V.

**TABLE S5**

Comparison of repeat content in grass genomes.

**TABLE S6**

Enrichment tests for increased repeat content and decreased GC content in windows flanking gaps.

**TABLE S7**

List of structural variants identified in *S. viridis*.

**TABLE S8**

Species used in OrthoFinder gene family analysis

**TABLE S9**

OrthoFinder gene counts per orthogroup

**TABLE S10**

OrthoFinder predicted duplications

**TABLE S11**

Gene Ontology enrichment analysis

**TABLE S12**

PlantiSMASH HMM enrichment analysis

## SUPPLEMENTARY DATASET LEGENDS

**SUPPLEMENTARY DATASET S1**

Chloroplast-derived contigs (FASTA).

**SUPPLEMENTARY DATASET S2**

ME034V gene Interproscan and PlantiSMASH functional annotations.

**SUPPLEMENTARY DATASET S3**

OrthoFinder gene families

**Figure S1.**
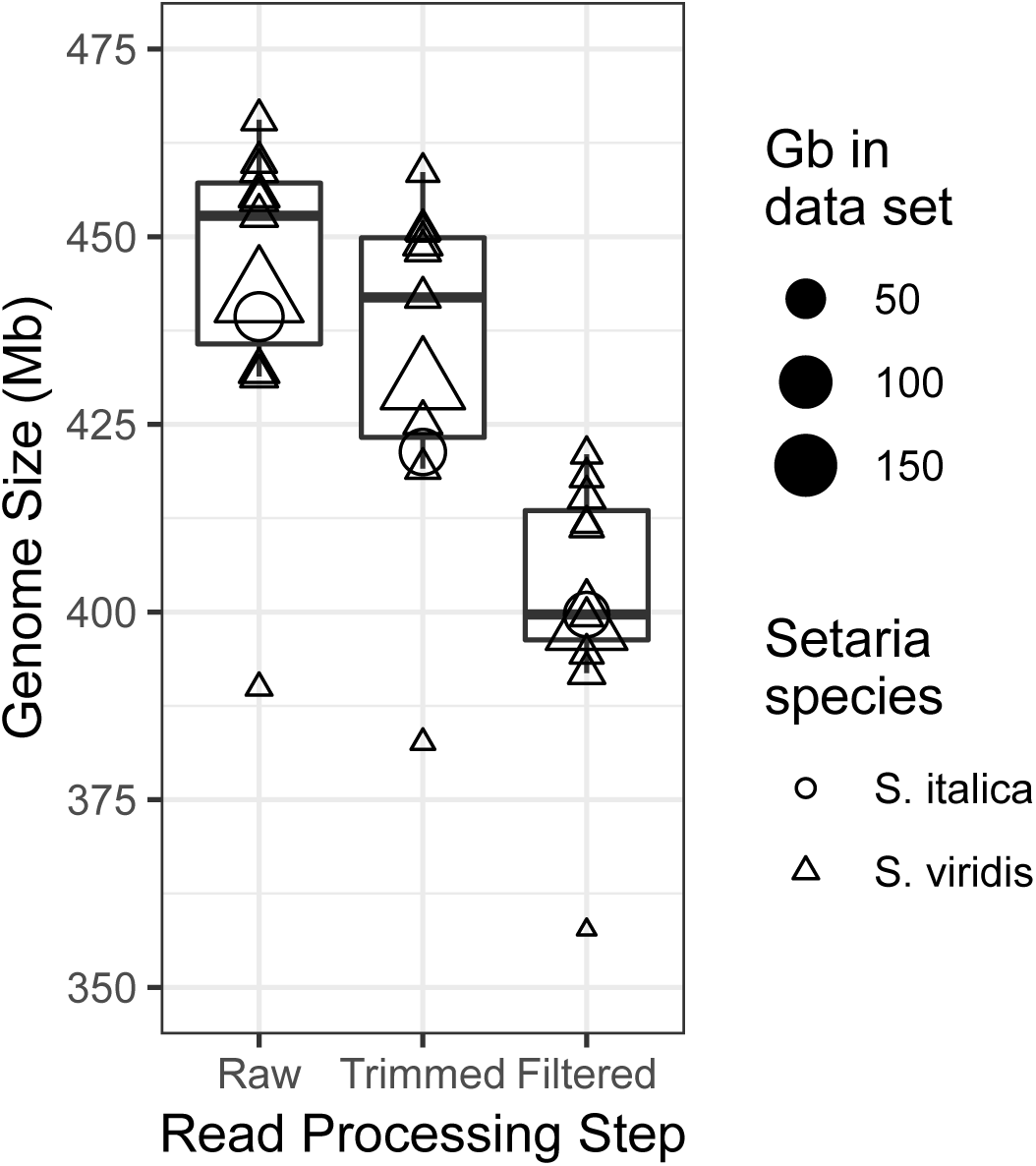
Estimated haploid genome length for *S. viridis* and *S. italica*. Data were evaluated without processing (Raw), after read quality trimming (Trimmed), and after quality trimming plus the removal of any reads that align to the chloroplast or mitochondria (Filtered). The amount of reads that remained following any processing is indicated by the relative size of the circles and triangles for cultivars of *S. italica* and *S. viridis*, respectively.

**Figure S2.**
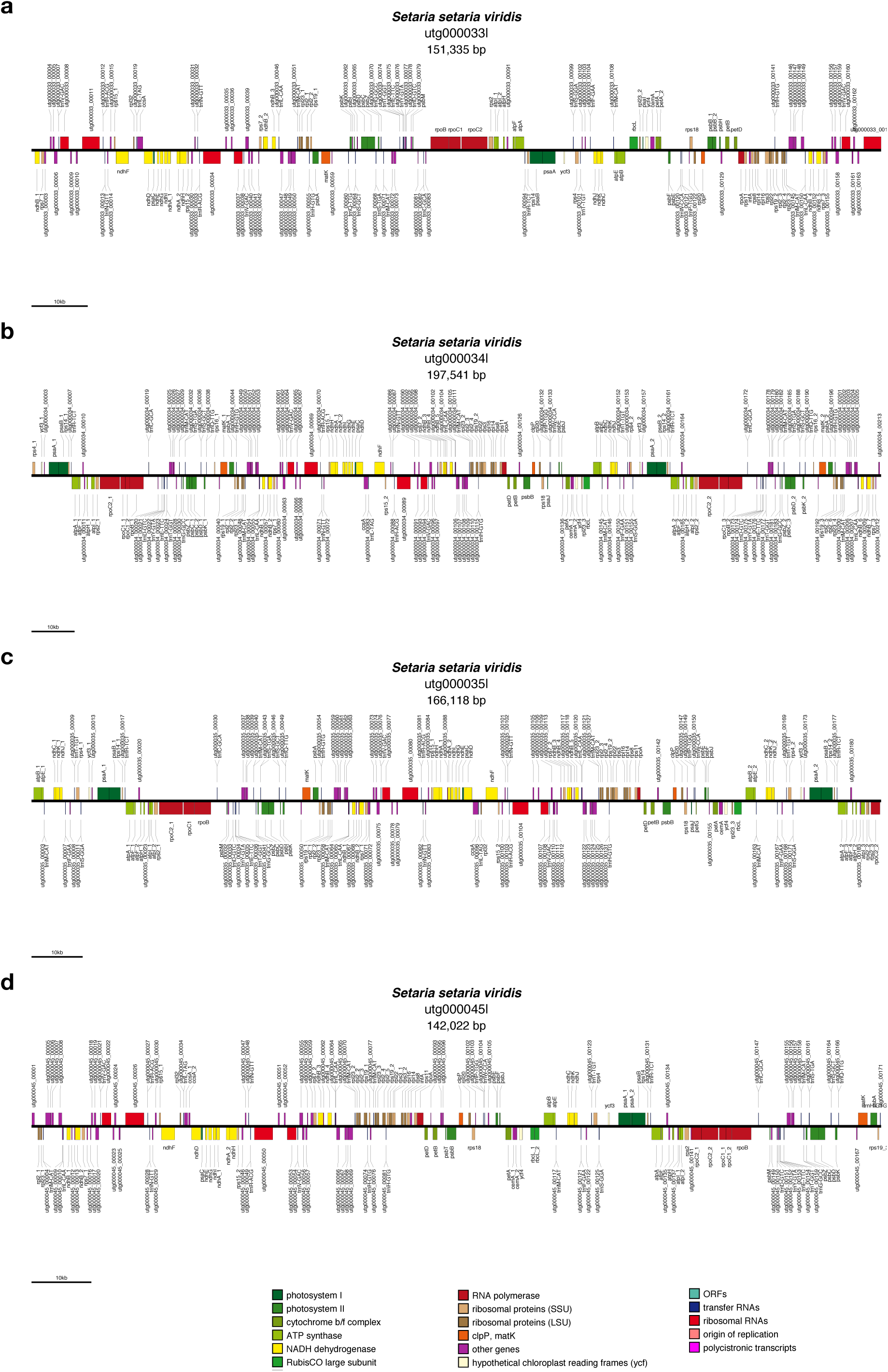
Physical maps of the four chloroplast-derived contigs. a) utg000033l, b) utg000034l, c) utg000035l, d) utg000045l.

**Figure S3.**
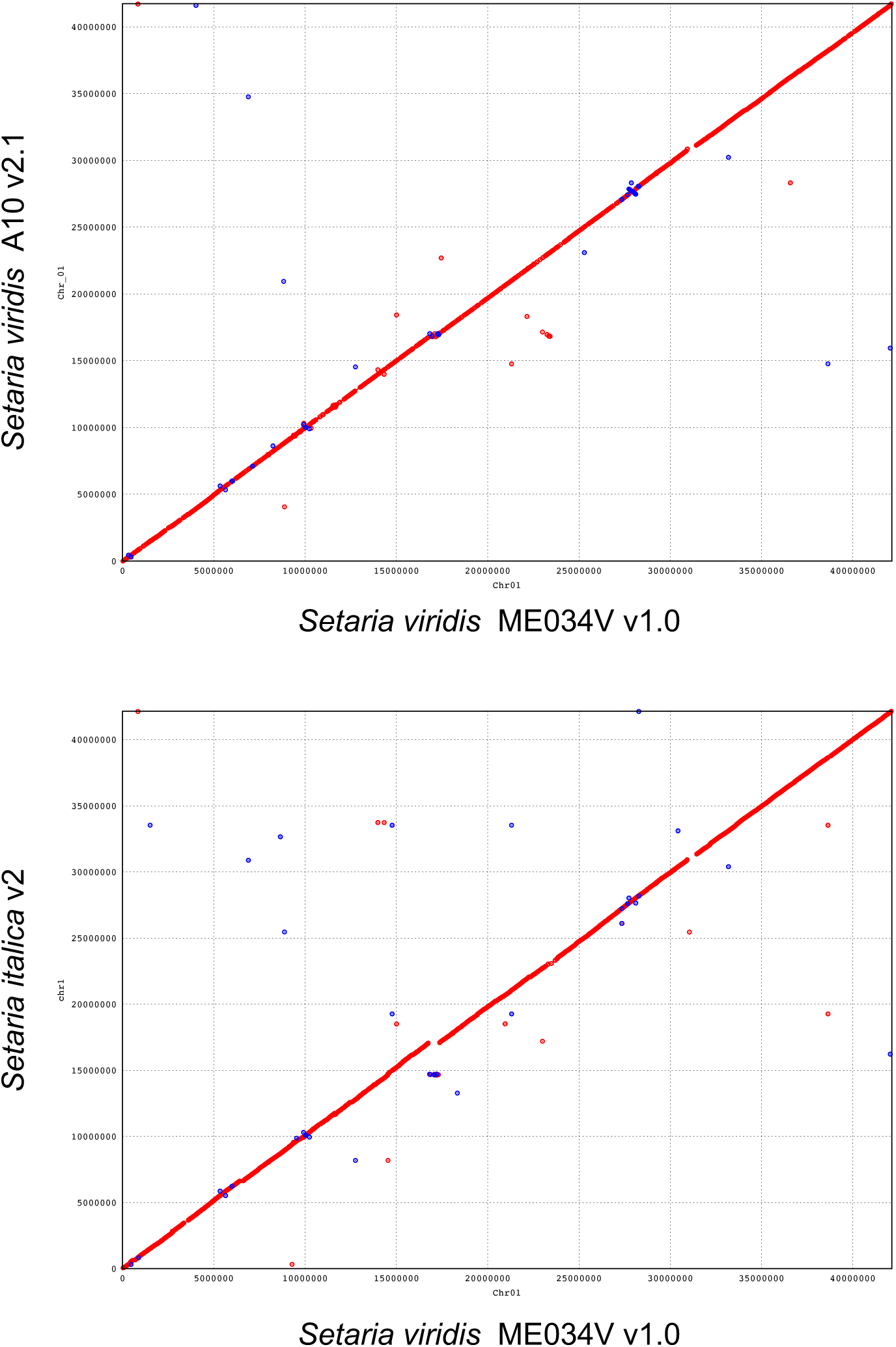
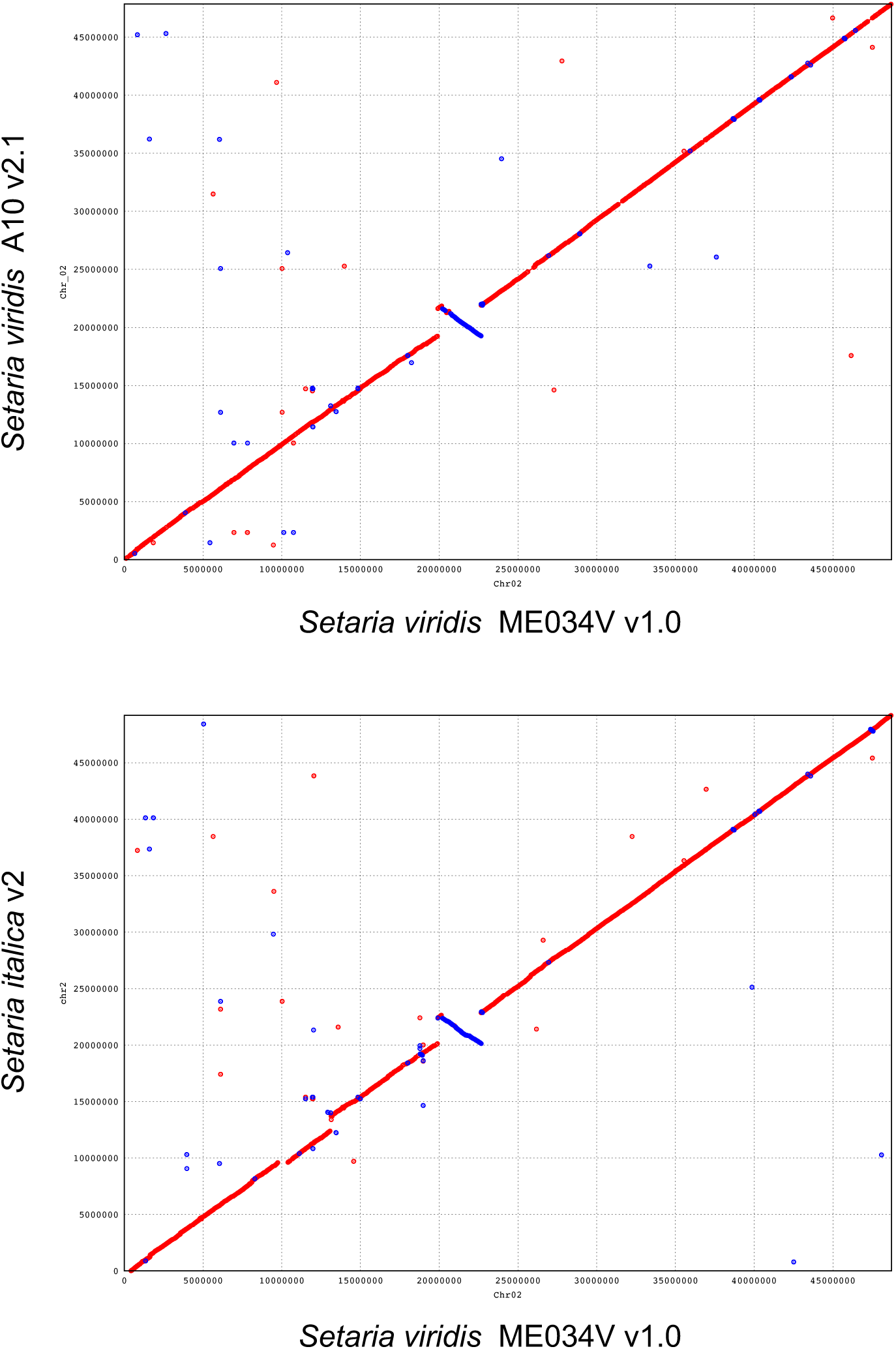
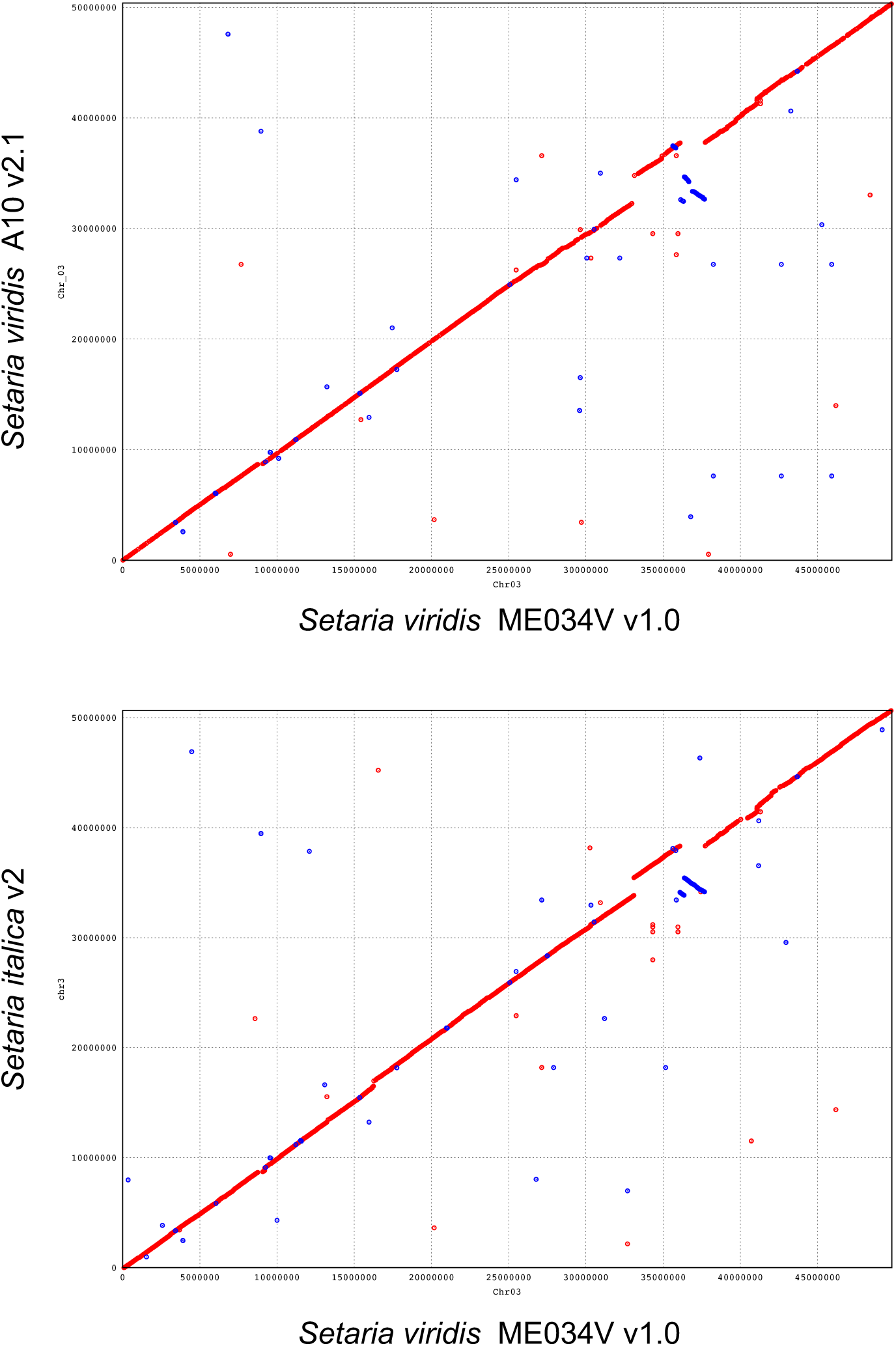
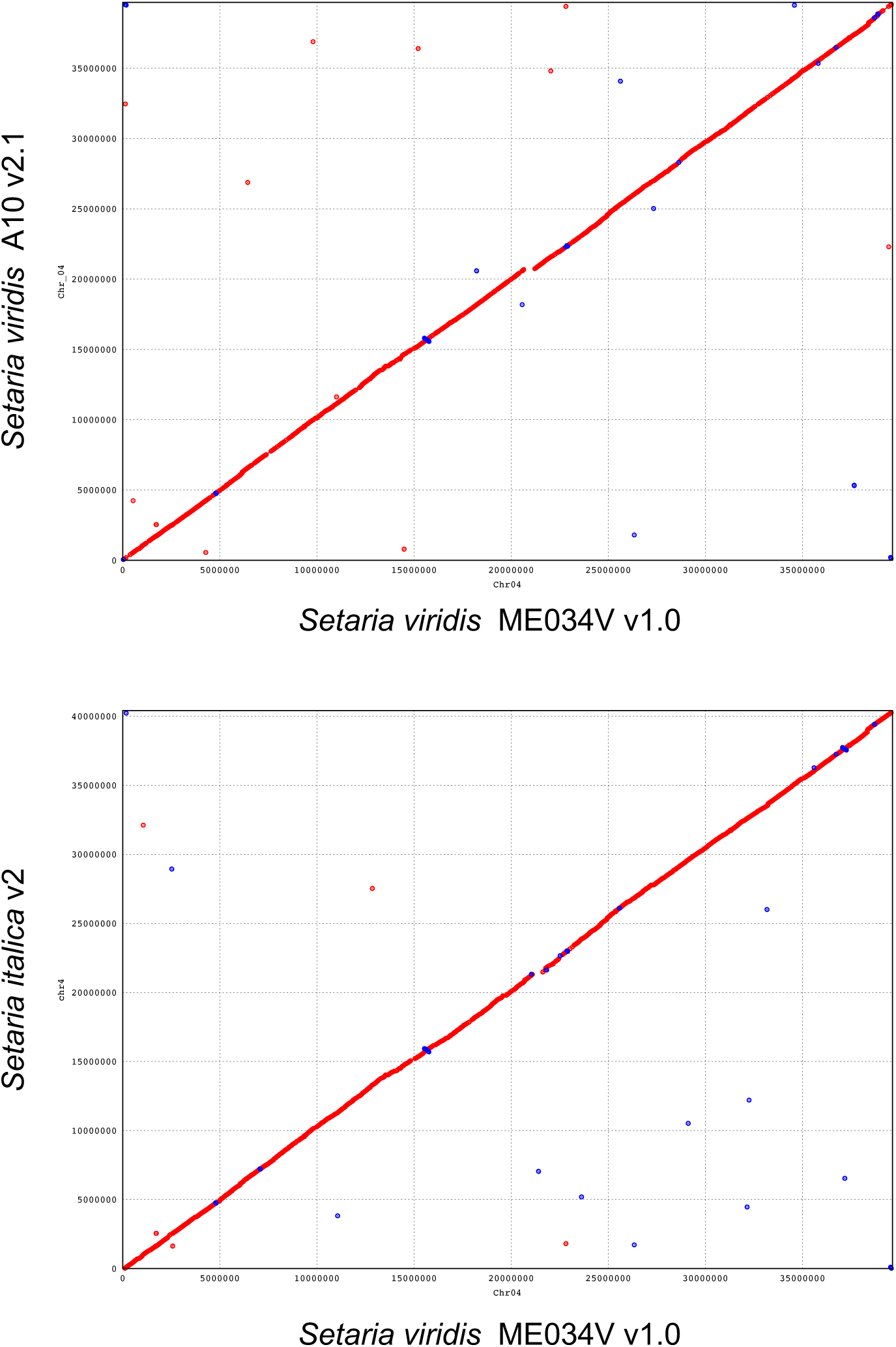
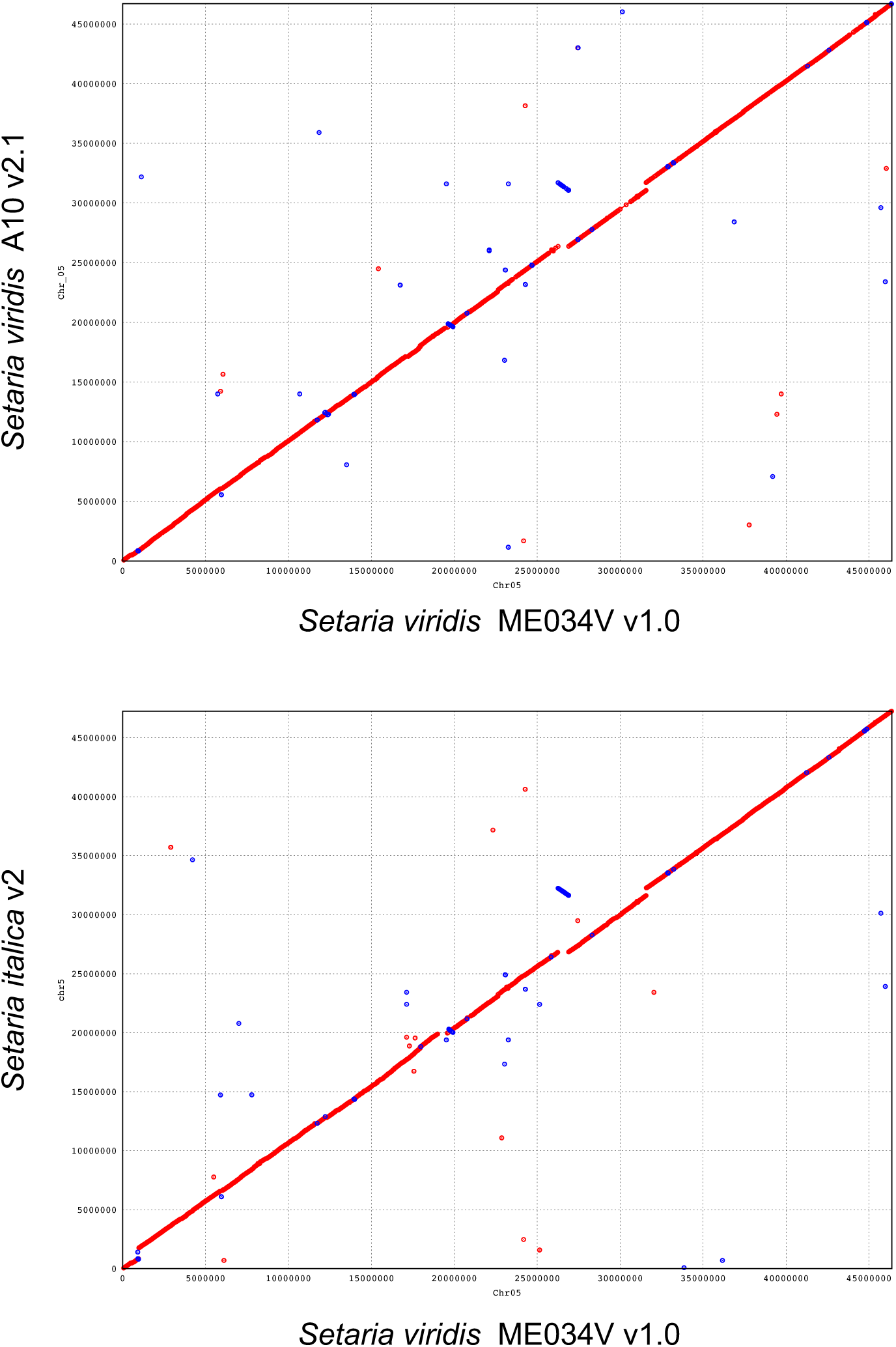
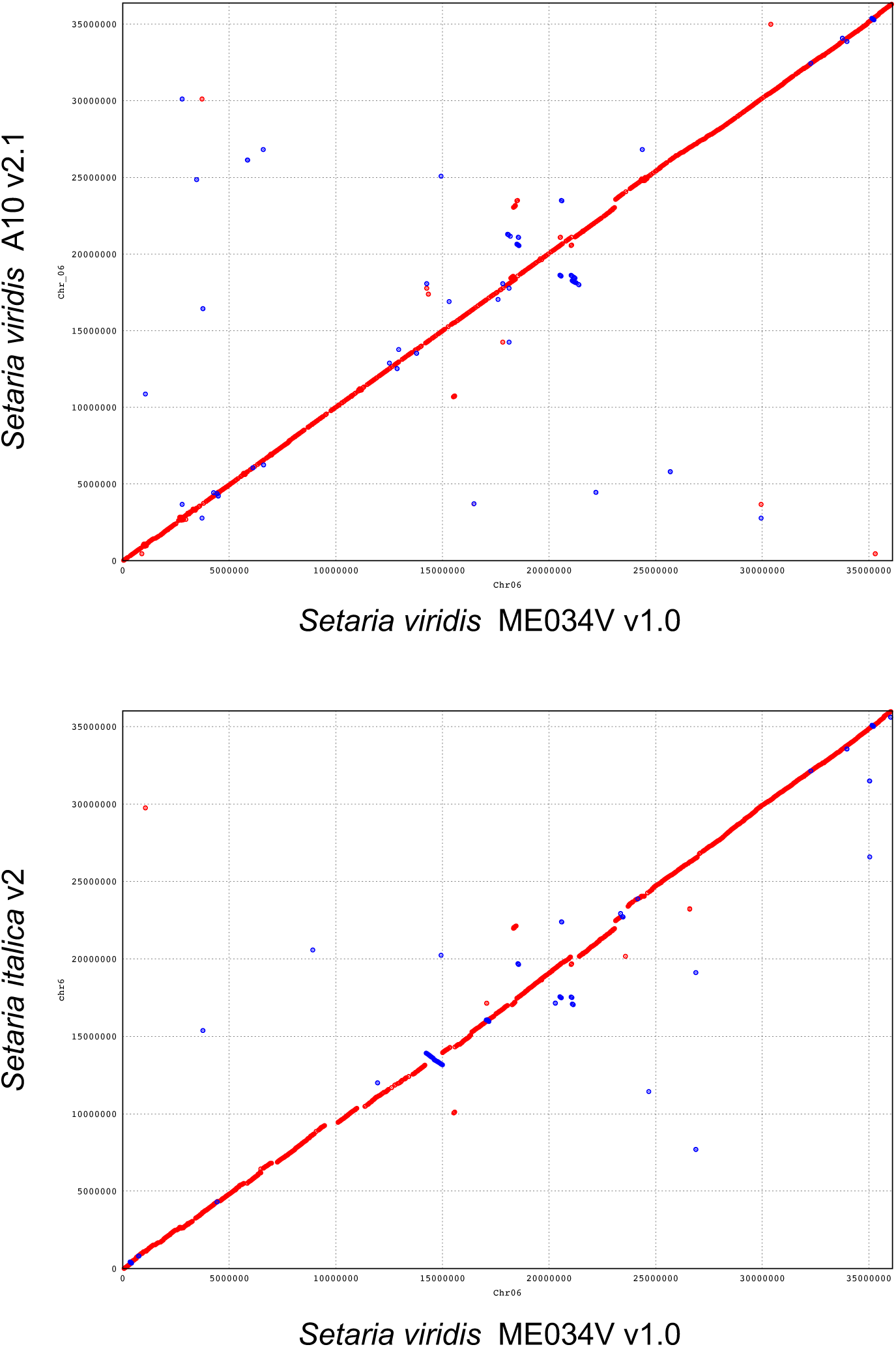
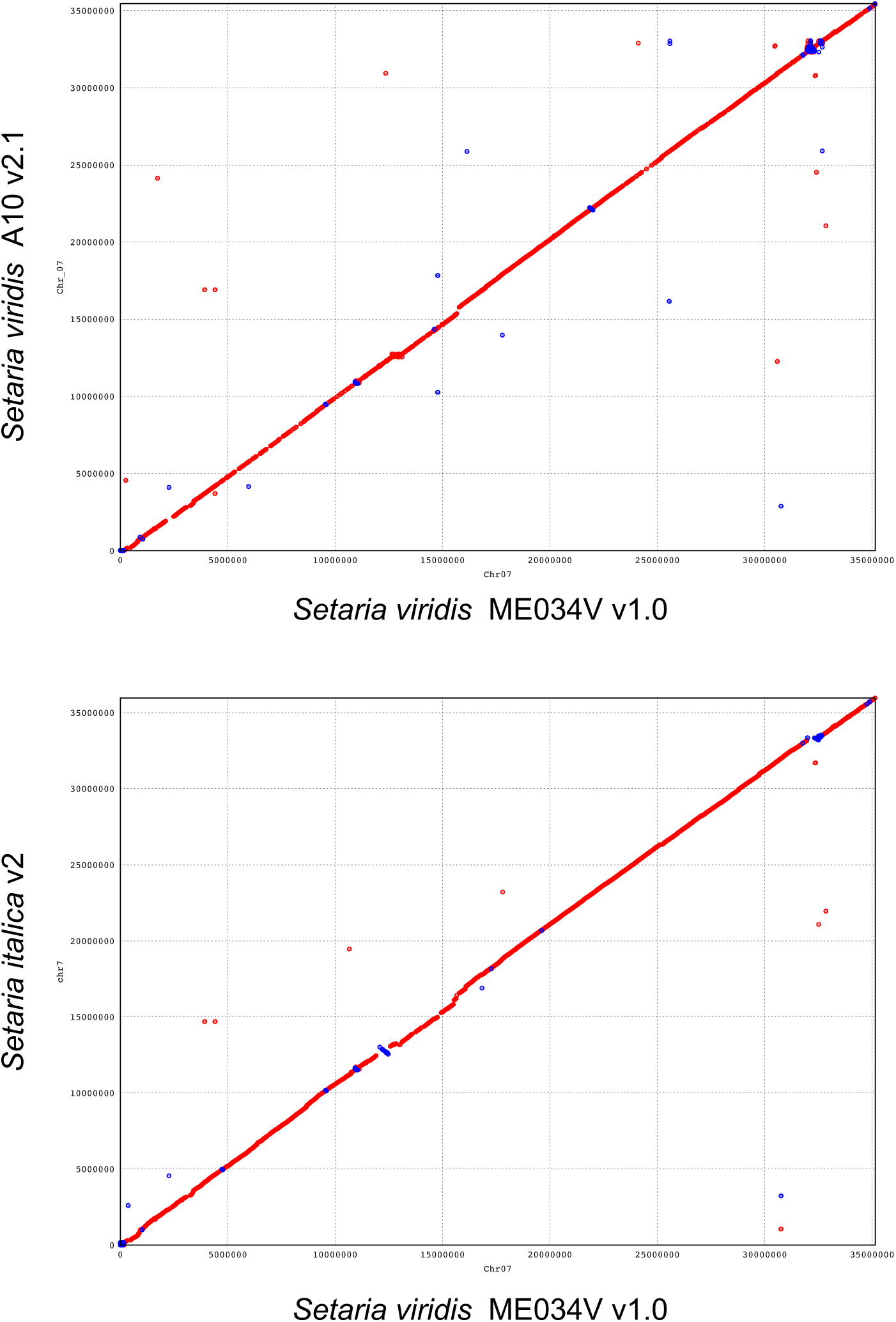
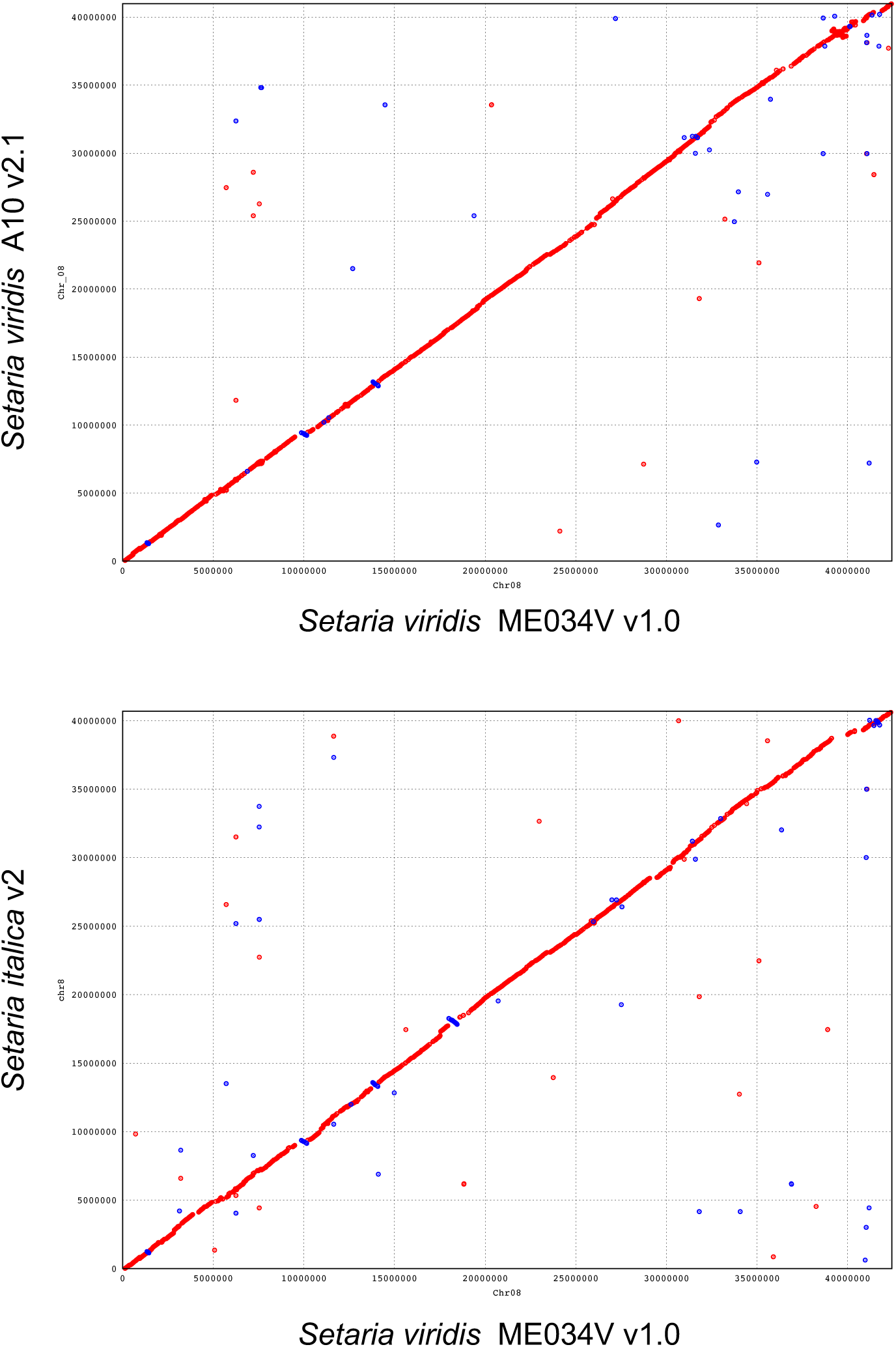
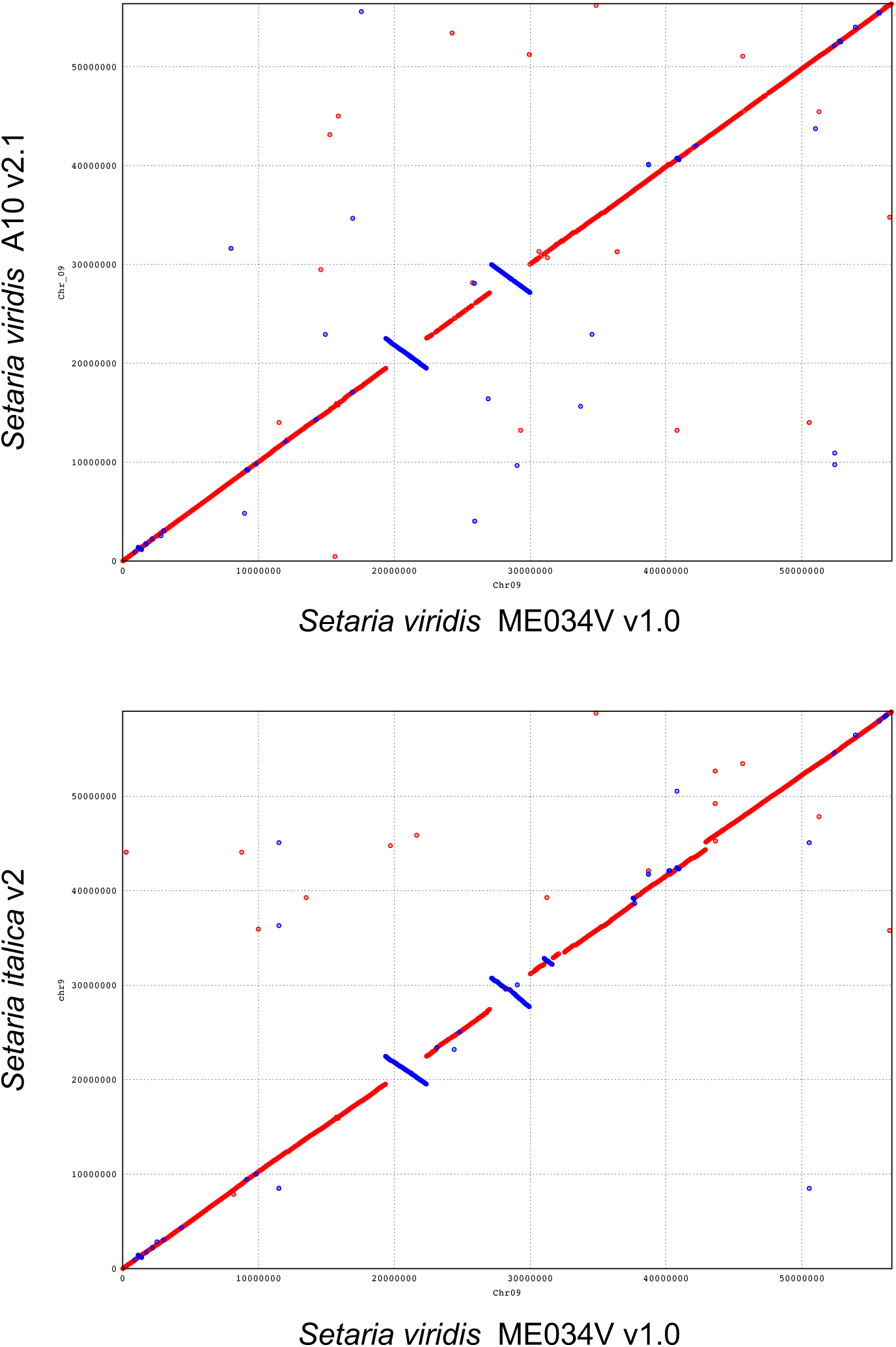
Chromosome alignments of ME034V to A10.1 and *S. italica*. a) Chromosome 1 b) Chromosome 2 c) Chromosome 3 d) Chromosome 4 e) Chromosome 5 f) Chromosome 6 g) Chromosome 7 h) Chromosome 8 i) Chromosome 9

**Figure S4.**
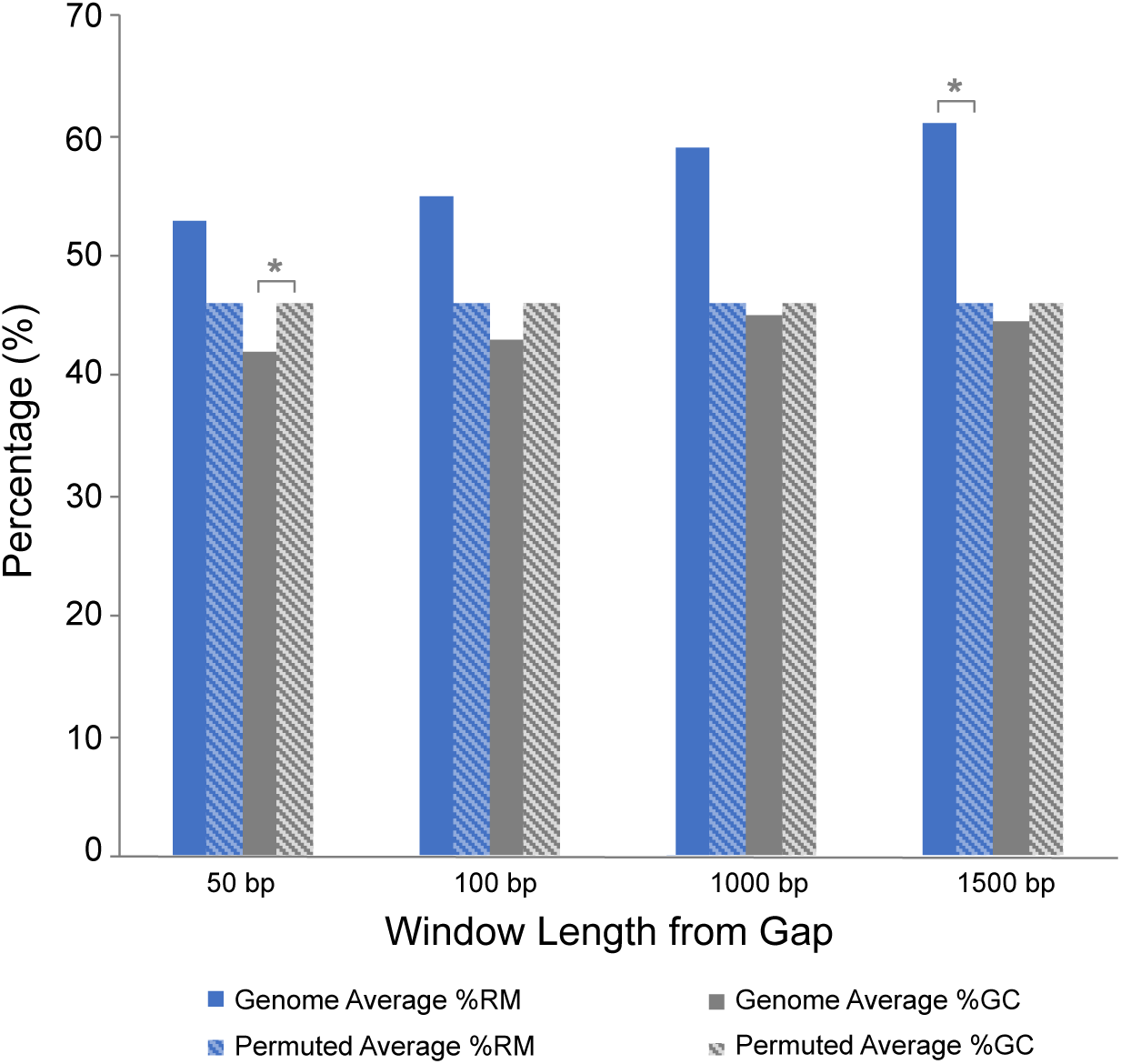
Statistical enrichment of repeat and low-GC near ME034V assembly gaps. One thousand randomized permutations of gap positions were generated to compare the true density of repeats and GC content with permuted values. The assessments were made with sequence from 50, 100, 1000, and 1500 bp from the gap flanks. P-values were calculated as the number of instances where the permuted values were either higher (for repeats) or lower (for %GC) than the observed values.

**Figure S5.**
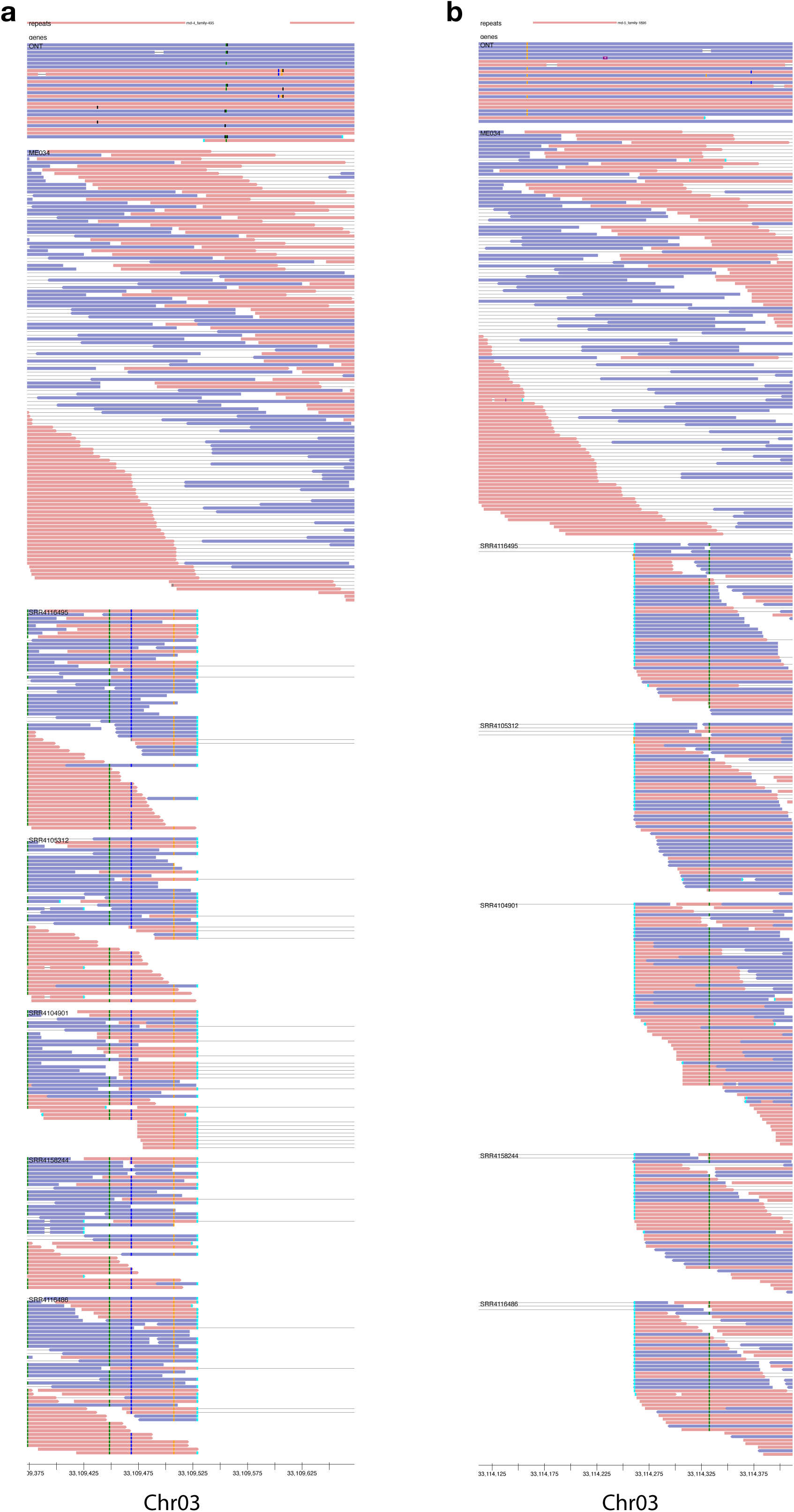
Read support for the *copia* Insertion (DEL0053315). A homozygous deletion (DEL00053315) was bioinformatically predicted in multiple *Setaria* samples which correspond to an LTR-rich locus around Chr03:33.2 Mb in ME034V. The left (a) and right (b) flanking regions illustrate alignments of ME034V Oxford Nanopore (ONT) reads, and paired Illumina reads from ME034V, SRR4116495, SRR4105312, SRR4104901 (Table S4). Pairs with too long of an insertion size (denoted by long gray bars connecting reads) and split reads (cyan box at read terminus) are indicated. SNPs are colored boxes of purple, green, dark blue, and orange while gaps are black.

**Figure S6.**
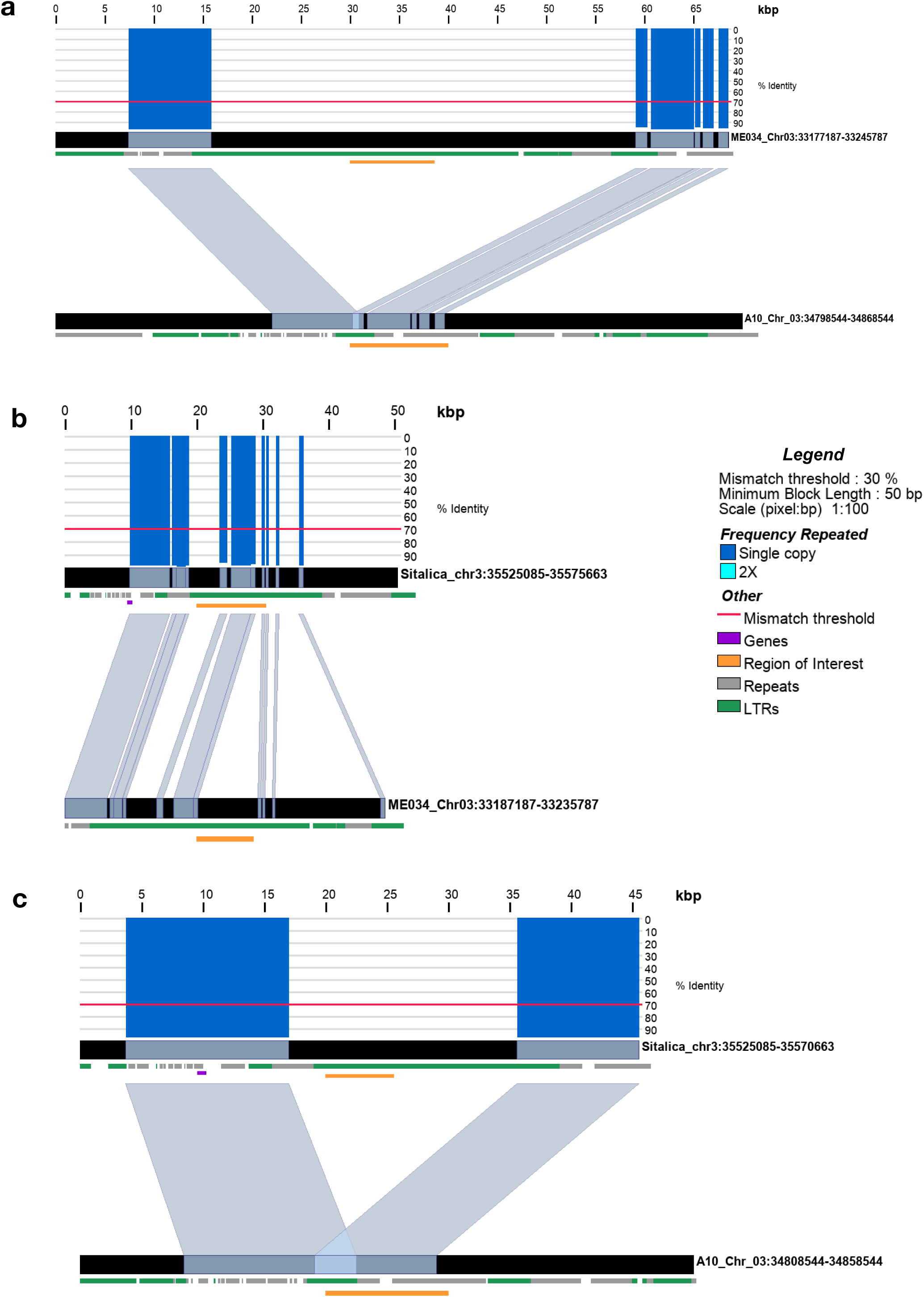
Genome view of *copia* insertion (DEL0053315). Synteny alignments between ME034V and A10 (a), ME034V and *S. italica* (b), and *S. italica* and A10 (c) showing the homozygous deletion (DEL00053315) of *copia-*rich region in A10. Blue-grey bars connect the two genomes when DNA sequence with >70% identity is observed (red line indicates threshold). Bars above the top track indicate sequence identity along the chromosomal segment from 0-100%, with the color indicating wither single copy (blue) or double copy (cyan) matches. Green tracks represent LTR elements while gray tracks are all other repeats repeat element classes. Orange tracks indicate the 1:1 homologous region to the predicted deletion site.

**Figure S7.**
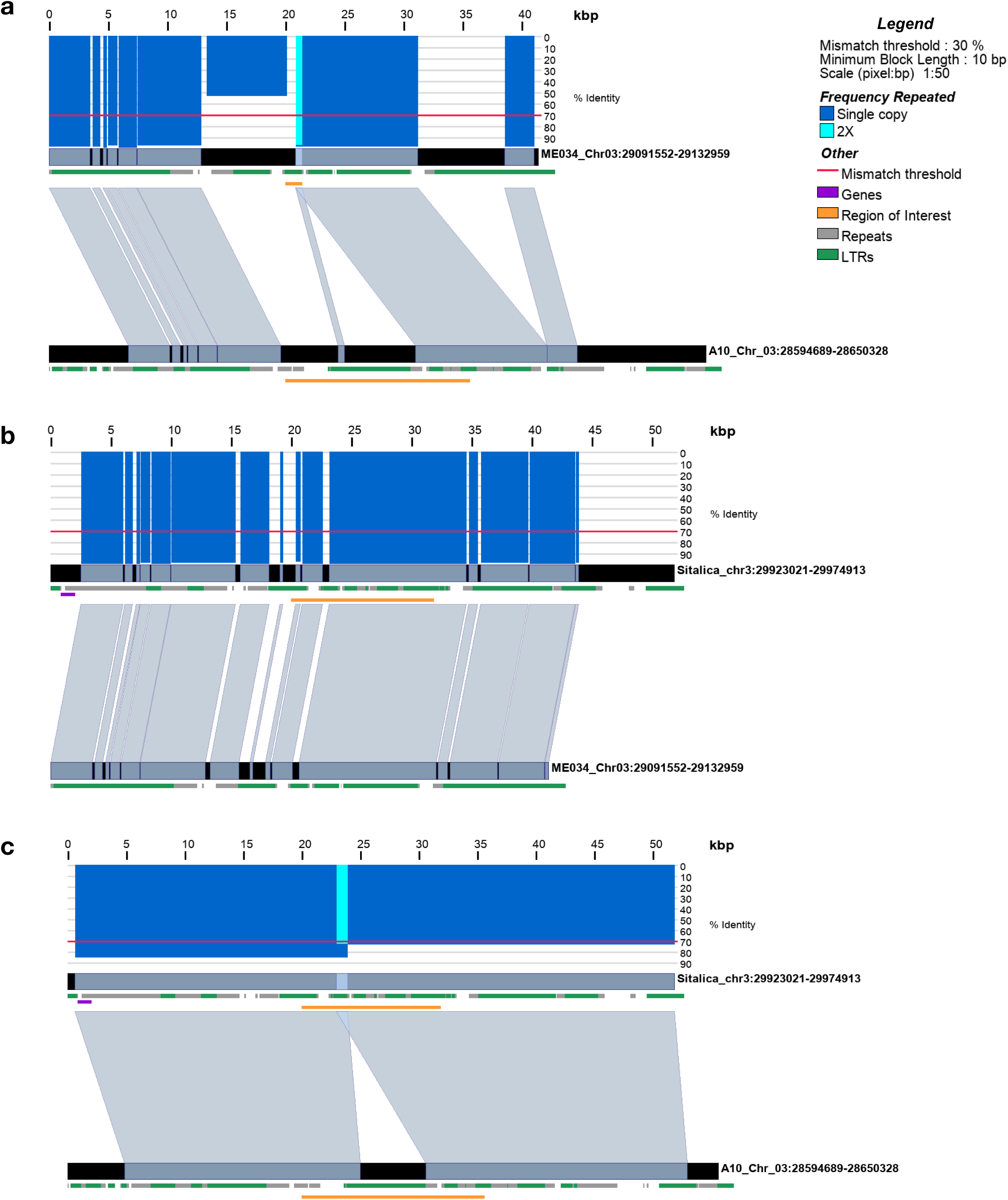
Genome view of *copia* insertion (DEL00051066). Synteny alignments between ME034V and A10 (a), ME034V and *S. italica* (b), and *S. italica* and A10 (c) showing a homozygous *copia* insertion unique to the A10 assembly. Blue-grey bars connect the two genomes when DNA sequence with >70% identity is observed (red line indicates threshold). Bars above the top track indicate sequence identity along the chromosomal segment from 0-100%, with the color indicating wither single copy (blue) or double copy (cyan) matches. Green tracks represent LTR elements while gray tracks are all other repeats repeat element classes. Orange tracks indicate the 1:1 homologous region to the predicted deletion site.

**Figure S8.**
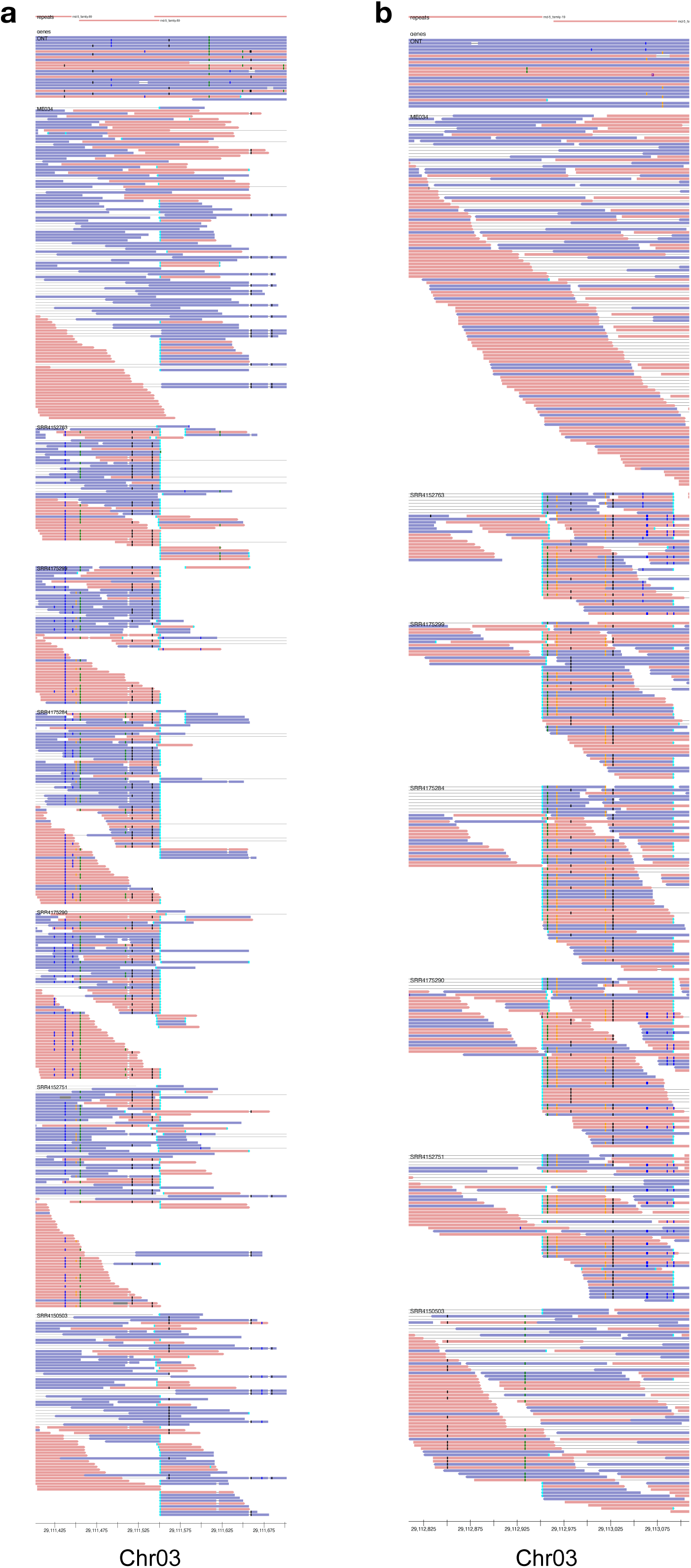
Read support for *copia* insertion (DEL00051066). A *copia* insertion in the A10 assembly relative to ME034V and *S. italica* was discovered after assessing the deletion locus DEL00051066. The insertion was identified in multiple *Setaria* samples around Chr03:29.1 Mb in ME034V. Left (a) and right (b) flanking regions illustrate alignments of ME034V ONT reads, ME034V paired Illumina reads, and other *Setaria* cultivars (see label on track for identifier; Table S4). Overlapping split paired reads, indicative of an insertion, are denoted with a bright blue box at the end of the read. SNPs are colored boxes of purple, green, dark blue, and orange while gaps are black.

**Figure S9.**
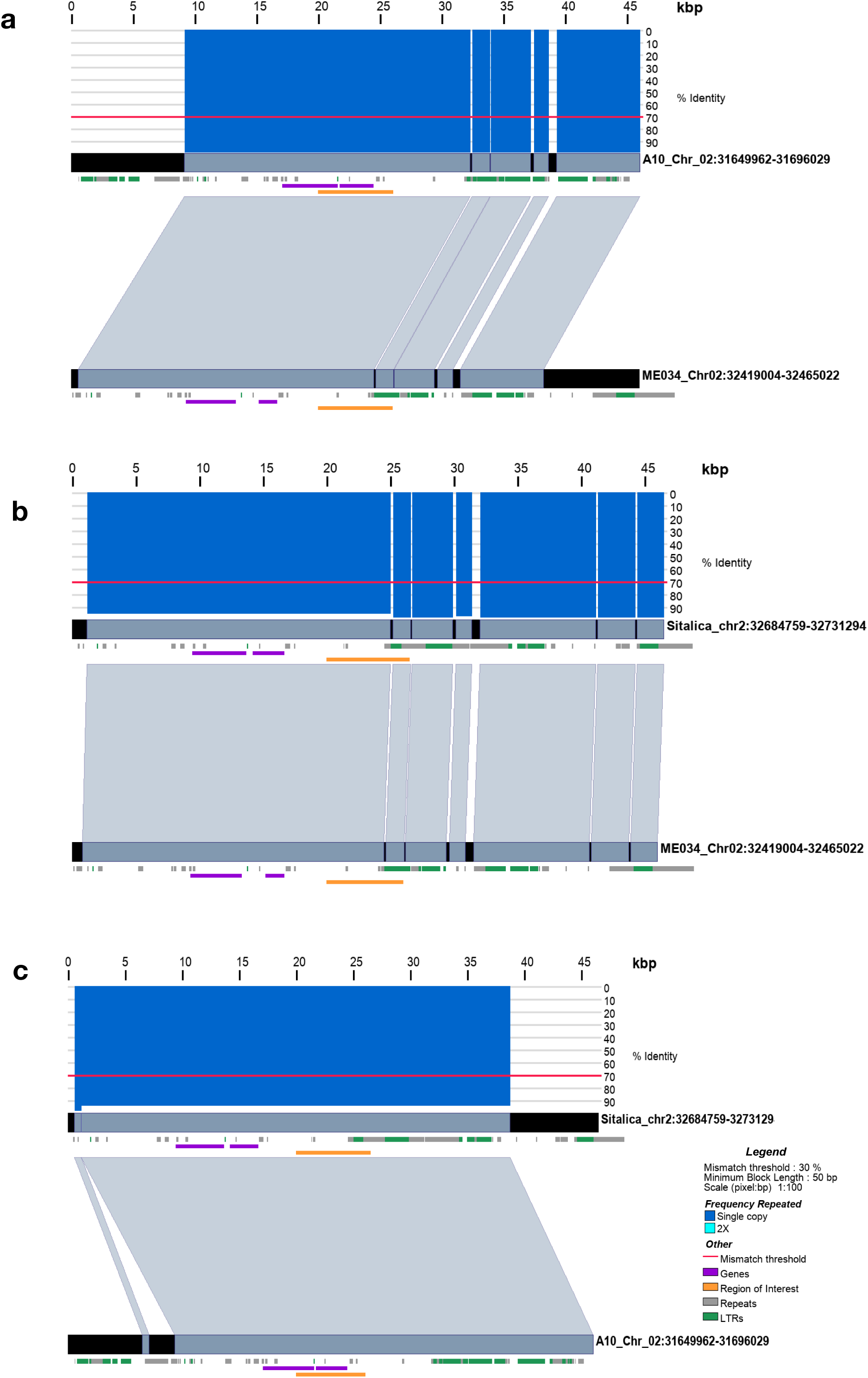
Genome view of gypsy insertion (DEL00033261). Synteny alignments between ME034V and A10 (a), ME034V and *S. italica* (b), and *S. italica* and A10 (c) showing the presence of the *gypsy* element in all three genomes. Blue-grey bars connect the two genomes when DNA sequence with >70% identity is observed (red line indicates threshold). Bars above the top track indicate sequence identity along the chromosomal segment from 0-100%, with the color indicating wither single copy (blue) or double copy (cyan) matches. Green tracks represent LTR elements while gray tracks are all other repeats repeat element classes. Orange tracks indicate the 1:1 homologous region to the predicted deletion site.

**Figure S10.**
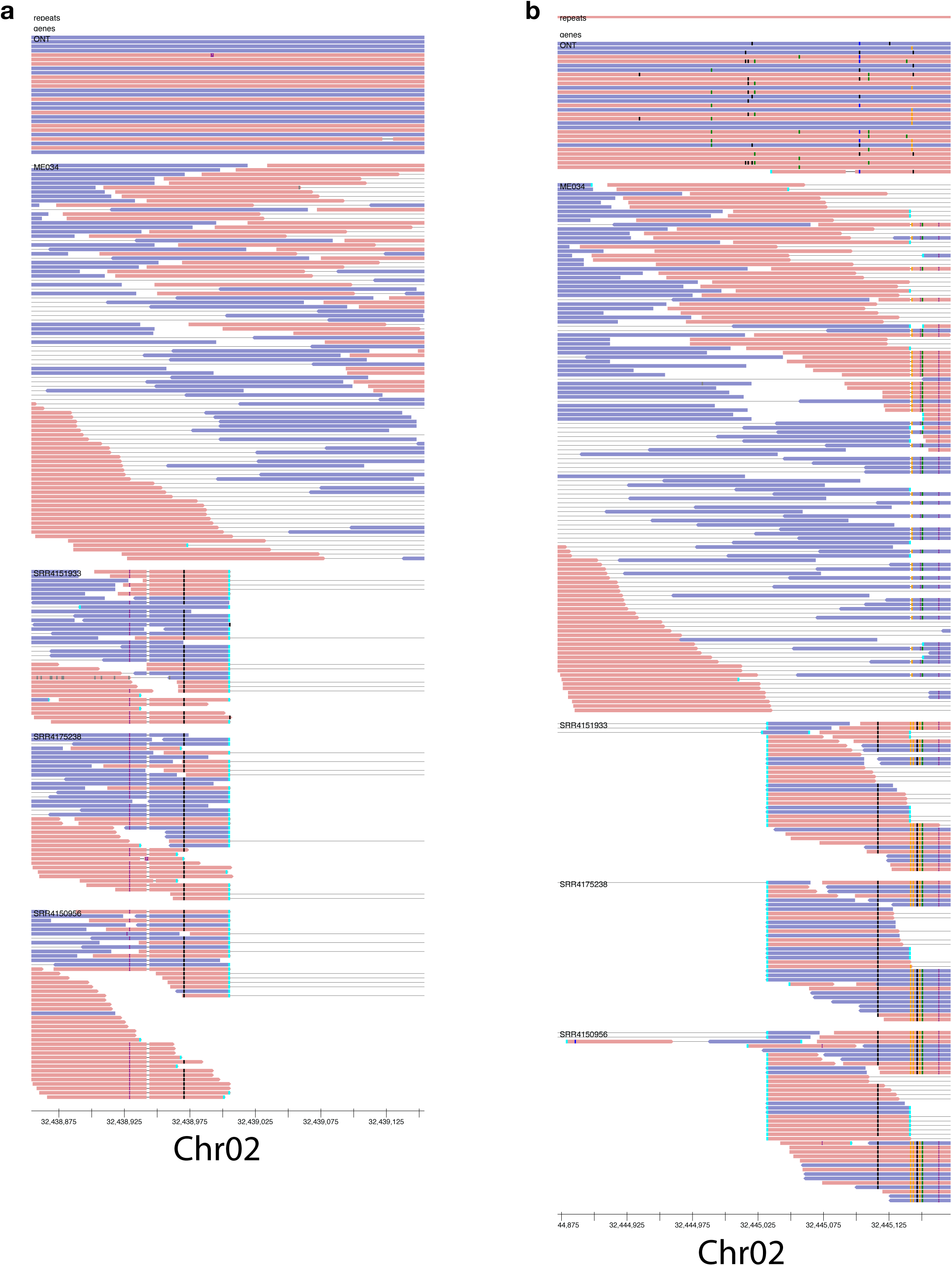
Read support for *gypsy* insertion (DEL00033261). A *gypsy* element present in all three *Setaria* assemblies (approximate location Chr02:32.4 Mb in ME034V) is missing in three other *S. viridis* samples SRR4151933 (cultivar Feldman_MF156), SRR4175238 (cultivar Estep_ME018), and SRR4150956 (cultivar Feldman_MF137). Left (a) and right (b) flanking regions illustrate alignments of ME034V ONT reads, ME034V paired Illumina reads, and other *Setaria* cultivars (Table S4). Pairs with too long of an insertion size (denoted by long gray bars connecting reads) and split reads (bright blue box at read terminus) are indicated. SNPs are colored boxes of purple, green, dark blue, and orange while gaps are black.

